# m^6^A epitranscriptomic modification regulates neural progenitor-to-glial cell transition in the retina

**DOI:** 10.1101/2022.05.08.491092

**Authors:** Yanling Xin, Qinghai He, Huilin Liang, Jingyi Guo, Qi Zhong, Ke Zhang, Jinyan Li, Yizhi Liu, Shuyi Chen

## Abstract

*N*^6^-methyladenosine (m^6^A) is the most prevalent mRNA internal modification and has been shown to regulate the development, physiology and pathology of various tissues. However, the functions of the m^6^A epitranscriptome in the visual system remain unclear. In this study, using a retina-specific conditional knockout mouse model, we show that retinas deficient in *Mettl3*, the core component of the m^6^A methyltransferase complex, exhibit structural and functional abnormalities beginning at the end of retinogenesis. Immunohistological and scRNA-seq analyses of retinogenesis processes reveal that retinal progenitor cells (RPCs) and Müller glial cells are the two cell types primarily affected by *Mettl3* deficiency. Integrative analyses of scRNA-seq and MeRIP-seq data suggest that m^6^A fine-tunes the transcriptomic transition from RPCs to Müller cells by promoting the degradation of RPC transcripts, the disruption of which leads to abnormalities in late retinogenesis and compromises the glial functions of Müller cells. Finally, overexpression of m^6^A-regulated RPC-enriched transcripts in late RPCs partially recapitulates the *Mettl3*-deficient retinal phenotype. Collectively, our study reveals an epitranscriptomic mechanism governing progenitor-to-glial cell transition during late retinogenesis, which is essential for the homeostasis of the mature retina. The mechanism revealed in this study might also apply to other nervous systems.

## Introduction

The retina is the neural sensory component of the visual system that is responsible for light perception and preliminary visual information processing. As a lateral derivative of the neural tube, the retina shares basic developmental, structural, and physiological principles with other neural tissues and has become an excellent model for studying neural biology. The retina is composed of six types of neurons, including rod and cone photoreceptors, bipolar cells, horizontal cells, amacrine cells, and retinal ganglion cells, that form complex neural circuits to detect and process visual signals (Masland, 2001). In addition to neurons, the retina contains one type of glial cell, Müller cells. As the sole glial cell inside the retina, Müller cells play pivotal roles in maintaining structural, physiological, and functional homeostasis in the neural retina (Newman and Reichenbach, 1996; Vecino et al., 2016). During development, all retinal neurons and Müller cells are derived from multipotent retinal progenitor cells (RPCs), which go through a series of competent states to give rise to various types of retinal cells in a sequential while overlapping manner. For example, in mice, retinal ganglion cells, cone photoreceptors, and GABAergic amacrine cells are born early before birth, while glycinergic amacrine cells, rod photoreceptors, and bipolar cells are born late during the postnatal period before RPCs finally become Müller glial cells (Agathocleous and Harris, 2009; Bassett and Wallace, 2012; Cepko, 2014; Heavner and Pevny, 2012). Similar to glial cells in other neural tissues, Müller cells maintain significant levels of gene expression signatures of progenitors (Blackshaw et al., 2004; Nelson et al., 2011; Roesch et al., 2008), which has encouraged people to explore the *in vivo* reprogramming potentials of Müller cells for treating retinal degeneration disease purposes in recent years (Hoang et al., 2020; Jorstad et al., 2017; Yao et al., 2018; Zhou et al., 2020). However, to function as glial cells, Müller cells must develop and maintain their unique transcriptome distinct from that of RPCs (Lin et al., 2019; Nelson et al., 2011). How the transcriptome of Müller cells is established and maintained is currently not well understood.

RNA modifications constitute an important layer of posttranscriptional regulation that orchestrates the metabolism, location and function of transcripts, which are collectively referred to as the epitranscriptome. Among the various modes of mRNA modification, *N*^6^-methyladenosine (m^6^A) is the most prevalent internal modification (Meyer and Jaffrey, 2014, 2017; Roundtree et al., 2017; Shi et al., 2019; Zaccara et al., 2019). m^6^A is cotranscriptionally installed on RNAs in the nucleus by a multisubunit methyltransferase ‘writer’ complex, in which METTL3 and METTL14 form the catalytic core; but only METTL3 has methyltransferase catalytic activity, while METTL14 functions as an allosteric activator (Bokar et al., 1994; Bokar et al., 1997; Liu et al., 2014; Sledz and Jinek, 2016; Wang et al., 2016a; Wang et al., 2016b). Two enzymes, FTO and ALKBH5, have been demonstrated to have m^6^A demethylase ‘eraser’ activity, emphasizing the reversible nature of the modification (Jia et al., 2011; Zheng et al., 2013). The effects of m^6^A on RNAs are mediated by various m^6^A readers, such as YTH family and HNRNP family proteins (Lee et al., 2020; Zaccara et al., 2019; Zhao et al., 2017). Gene knockdown and genetic knockout studies on m^6^A writers, erasers and readers have shown that m^6^A plays important roles in regulating various aspects of organism development, physiology, and disease progression (Frye and Blanco, 2016; Liu et al., 2019; Livneh et al., 2020). However, the functions of the m^6^A epitranscriptome in the visual system remain unclear.

In this study, using a mouse model of retina-specific conditional knockout of *Mettl3* and combining scRNA-seq, MeRIP-seq and gene expression manipulation, we reveal that the m^6^A epitranscriptome plays critical roles during late retinogenesis by fine-tuning the transcriptomic transition from RPCs to Müller cells, which is essential for the structural and physiological homeostasis of the mature retina.

## Results

### *Mettl3* deficiency leads to structural and physiological abnormalities in the retina

To investigate the function of the m^6^A epitranscriptome in the retina, we conditionally knocked out *Mettl3*, the core component of the m^6^A-writer complex, in RPCs using *Six3-Cre* (Furuta et al., 2000) and *Mettl3^floxed^* (Lin et al., 2017) mice. METTL3 is abundantly and ubiquitously expressed in the retina from early embryonic age to adult stage (Figure1-figure supplement 1A-1D). Immunofluorescence staining (IF) demonstrated that *Mettl3* was efficiently knocked out in the central retinas of *Six3-Cre^+^; Mettl3^floxed/floxed^* mice (hereafter referred to as *Mettl3^CKO^*) (Figure 1A and 1A’), and MeRIP-qPCR examination of several candidate genes showed that m^6^A levels were efficiently downregulated in *Mettl3^CKO^* retinas (Figure1-figure supplement 1E). Even though *Mettl3* was efficiently deleted from RPCs, the retinas of *Mettl3^CKO^* mice at p0 appeared grossly normal, exhibiting an already separated retinal ganglion cell layer (RGL) and a still-developing retinoblast layer (RBL), comparable to littermate control retinas (Figure 1B and 1B’). However, at p14, the time point when retinogenesis should have completed and mature retinal organization should have been established, histological examination showed that the *Mettl3^CKO^* retinas exhibited severe structural disorganization (Figure 1C and 1C’). At this age, the control retinas were well laminated into three cellular layers and two synaptic layers (Figure 1C). In the mutant retinas, although all the cellular and synaptic layers were distinguishable, the outer nuclear layer (ONL) presented many rosettes, and some inner nuclear layer (INL) cells were drawn into the ONL (Figure1C’, the asterisks indicate the rosette structures). We then examined the visual function of the retina using multifocal electroretinogram (mfERG). The results showed that the visual responses from most regions of *Mettl3^CKO^* retinas exhibited abnormal wave forms and much reduced P1 peak amplitudes compared with control retinas (Figure 1D), demonstrating disrupted visual function of the mutant retinas. To examine whether retinal neurons and glial cells were generated in structurally distorted *Mettl3^CKO^* retinas, we performed IF analyses of characteristic markers of various types of retinal cells. The IF results showed that all types of retinal cells, including RECOVERIN^+^ photoreceptors, VSX2^+^ bipolar cells, SOX9^+^ Müller glia, CALBINDIN^+^ horizontal cells, HPC-1^+^ amacrine cells, and BRN3A^+^ retinal ganglion cells, were generated in *Mettl3^CKO^* retinas, although some cells in the INL, especially bipolar cells, Müller glia, and horizontal cells, were displaced into the ONL (Figure1-figure supplement 2).

**Fig. 1.**
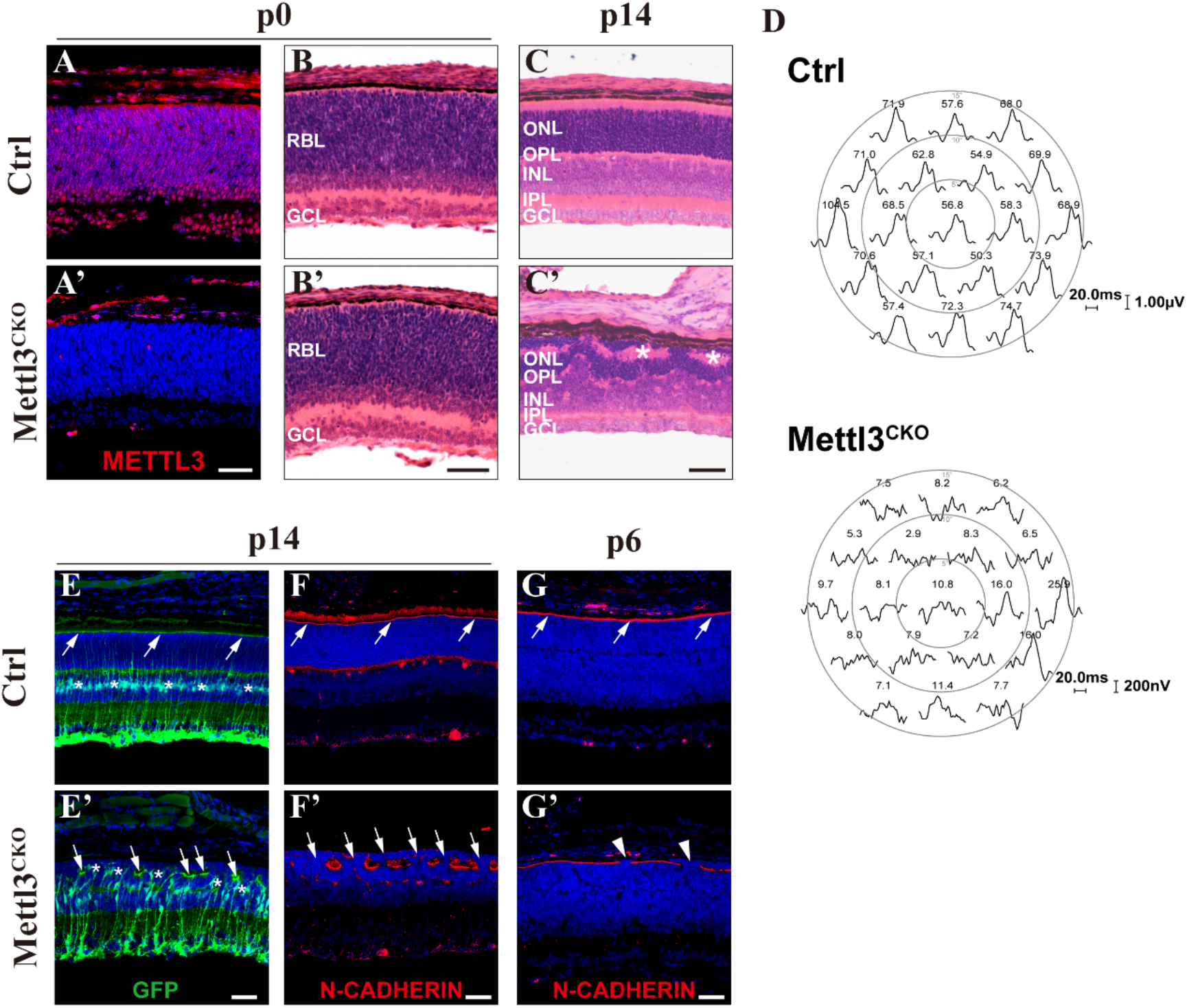
*Mettl3* deficiency leads to structural and physiological abnormalities in the retina. (A, A’) Confocal images of METTL3 expression in p0 control (Ctrl) and *Mettl3^CKO^* retinas showing that *Mettl3* was efficiently deleted in the mutant retinas. (B, B’) H&E staining of p0 control and *Mettl3^CKO^* retinas showing that the mutant retinas had a grossly normal histological structure. (C, C’) H&E staining of p14 control and *Mettl3^CKO^* retinas showing that the mutant retinal structure was severely disrupted. * indicates rosette structures. (D) Multifocal ERG response recordings in each hexagonal region of the retinas stimulated with 19 hexagonal light signals showing that the visual function of *Mettl3^CKO^* retinas was severely disrupted. The numbers on the top of each waveform plot indicate the averaged ERG response in the respective hexagonal area in nV/deg^2^. (E, E’) Confocal images of p14 retinas labeled with GFP to highlight Müller cells (mice on the *Rlbp1-GFP* background). The white arrows indicate the outer limiting membrane (OLM). * in E’ indicates mislocalized somata of Müller cells. (F, F’) Confocal images of p14 retinas stained for N-CADHERIN to highlight the OLM. The white arrows in F indicate the OLM, and those in F’ indicate breaks between the OLM. (G, G’) Confocal images of p6 retinas stained for N-CADHERIN showing that breaks in the OLM (white arrowheads) in *Mettl3^CKO^* retinas started to appear at p6. RBL: retinoblast layer; GCL: ganglion cell layer; ONL: outer nuclear layer; OPL: outer plexiform layer; INL: inner nuclear layer; IPL: inner plexiform layer. Scale bars are 50 μm.

In the retina, Müller cells are the primary glial cells that execute the glial supporting roles in maintaining the structural and functional homeostasis in the retina. Müller cells monitor the retina by extending elaborate cellular processes to interact with every retinal neuron and form an outer limiting membrane (OLM) and an inner limiting membrane (ILM) to ensheath the entire retina (Wang et al., 2017). Mechanical and genetic ablation studies have shown that loss of Müller cells leads to severe structural disorganization of the retina (Byrne et al., 2013; MacDonald et al., 2015; Shen et al., 2012), phenotypes similar to those observed in *Mettl3^CKO^* retinas. To examine how Müller cells were affected by *Mettl3* knockout, we placed *Mettl3^CKO^* mice on the *Rlbp1-GFP* mouse background, whose Müller glia express GFP (Vazquez-Chona et al., 2009). GFP IF illustrated severely distorted morphology of Müller glia in *Mettl3^CKO^* retinas: somata of many mutant Müller glia translocated from their normal position in the middle of the INL to move upward and mingle with photoreceptors in the ONL (Figure 1E’, the asterisks indicate the soma of Müller glia). Moreover, the apical edges of the mutant Müller glia failed to reach the apical edge of the retina, where they normally organize into the OLM; instead, they formed a discontinuous OLM that was separated into each rosette structure (Figure 1E’, the white arrows indicate the broken OLM). We then further examined the OLM by performing N-cadherin IF, which showed that the OLM in *Mettl3^CKO^* retinas became discontinuous and exhibited many breaks, through which many photoreceptors fled into the subretinal space (Figure 1F’, the white arrows indicate breaks in the OLM). By examining the retinas of pups at different postnatal ages, we found that occasional breaks along the OLM started to appear as early as p6 (Figure 1G’, the white arrowheads indicate breaks in the OLM), coinciding with the peak time point at which Müller glia are generated.

Taken together, these results demonstrate that *Mettl3* deficiency leads to structural and functional abnormalities in the mature retina and suggest that disfunction of Müller glial cells is a primary cause for these abnormalities.

### *Mettl3* deficiency distorts late-stage retinogenesis

We next performed BrdU labeling assays to examine the proliferative activity of RPCs. At E15.5, there were similar numbers of BrdU^+^ cells in *Mettl3^CKO^* retinas and control retinas (Figure2-figure supplement 1), suggesting that *Mettl3* deficiency does not affect the proliferation of early RPCs. However, at p1, fewer BrdU^+^ cells were observed in *Mettl3^CKO^* retinas than in control retinas (Figure 2A, 2A’ and2B), indicating that *Mettl3^CKO^* late RPCs proliferated slower than control RPCs. To examine whether the cell cycle progression of late RPCs was affected by *Mettl3* deletion, we harvested p1 retinas at 6 h after BrdU injection and subjected them to costaining with BrdU and pH3. We calculated the percentage of BrdU^+^; pH3^+^ RPCs vs. pH3^+^ RPCs, which showed that the percentage was significantly lower in *Mettl3^CKO^* retinas than in control retinas (Figure 2C, 2C’ and 2D), suggesting that S- to-M phase progression was prolonged in *Mettl3^CKO^* late RPCs. The postnatal period occurs at the end of retinogenesis, when an increasing number of RPCs withdraw from the cell cycle and undergo terminal differentiation. We examined the rate of cell cycle withdrawal of RPCs by injecting BrdU at p1 and monitored their KI67 expression 48 h later at p3. The results showed that there were significantly fewer BrdU^+^; KI67^-^ RPCs in *Mettl3^CKO^* retinas at p3 (Figure 2E, 2E’ and 2F), meaning that *Mettl3^CKO^* late RPCs withdrew from the cell cycle slower than control cells. Consistent with this slower withdrawal rate, at p5, when proliferating retinogenesis events were mostly terminated in control central retinas (Figure 2G, and 2H), a significant number of proliferating cells were still observable in *Mettl3^CKO^* central retinas (Figure 2G’, 2H). Taken together, these results demonstrate that the cell cycle progression of late RPC is distorted and retinogenesis is extended in *Mettl3^CKO^* retinas.

**Fig. 2.**
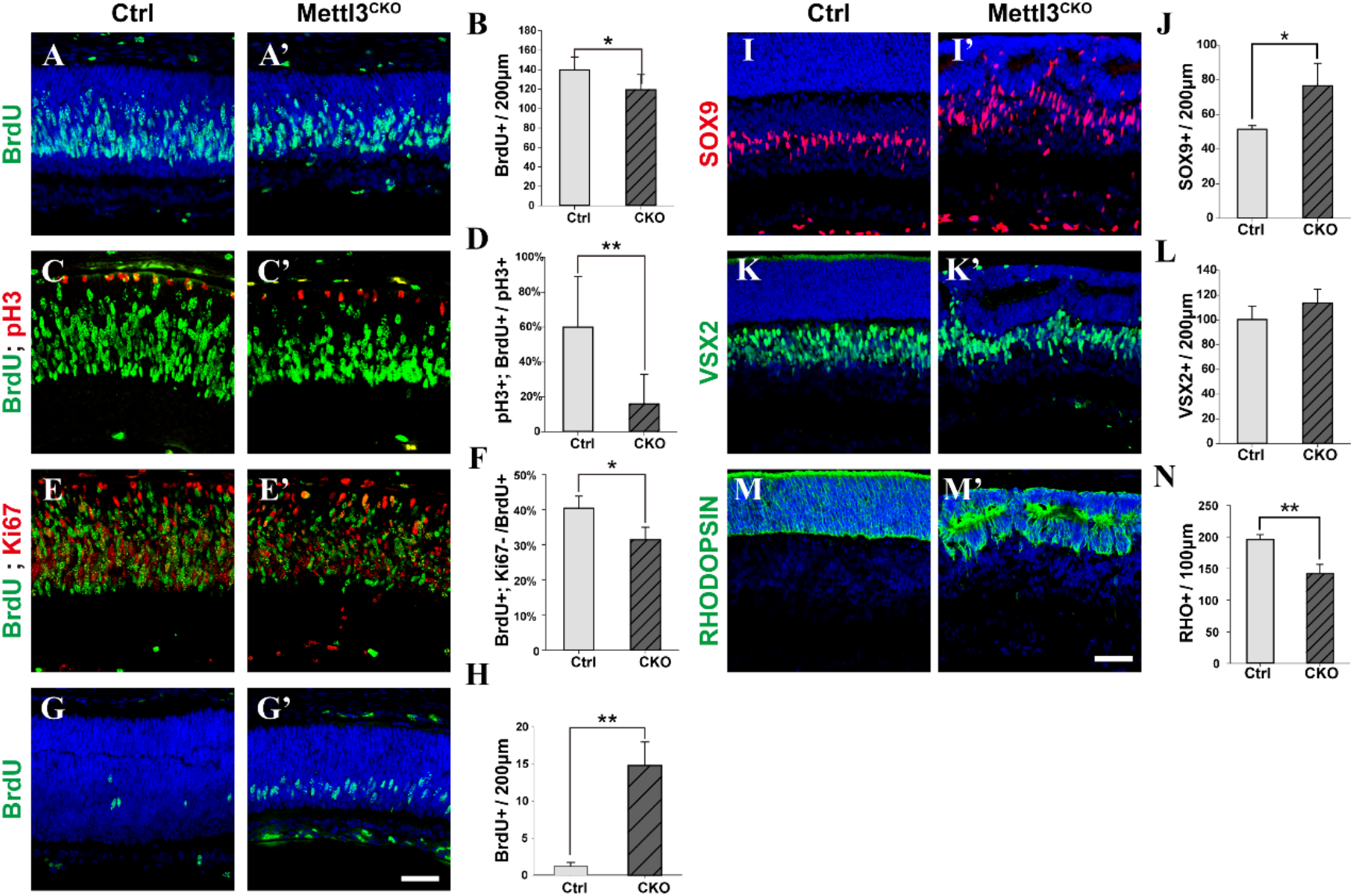
*Mettl3* deficiency distorts late-stage retinogenesis. (A, A’) Confocal images of p1 retinas stained for BrdU. The pups were injected with BrdU, and their retinas were collected 2 h later. (B) Quantification of the BrdU^+^ cells in A and A’. (C, C’) Confocal images of p1 retinas stained for BrdU and pH3. The pups were injected with BrdU, and their retinas were collected 6 h later. (D) Quantification of the BrdU^+^ and pH3^+^ cells in C and C’. (E, E’) Confocal images of p3 retinas stained for BrdU and KI67. The pups were injected with BrdU at p1, and their retinas were collected 48 h later. (F) Quantification of the BrdU^+^ and Ki67^-^ cells in E and E’. (G, G’) Confocal images of p5 retinas stained for BrdU. The pups were injected with BrdU, and their retinas were collected 2 h later. (H) Quantification of the BrdU^+^ cells in G and G’. (I, I’) Confocal images of p7 retinas stained for SOX9 to illustrate Müller cells. (J) Quantification of the SOX9^+^ cells in I and I’. (K, K’) Confocal images of p7 retinas stained for VSX2 to illustrate bipolar cells. (L) Quantification of the VSX2^+^ cells in K and K’. (M, M’) Confocal images of p7 retinas stained for RHODOPSIN to illustrate rods. (N) Quantification of the RHODOPSIN^+^ cells in M and M’. The data in B, D, F, H, J, L, and N are presented as the means ± standard deviations, corresponding to three independent biological replicates. * p < 0.05, ** p < 0.01. The scale bars in G’ and M’ are 50 μm and apply to A-G’ and I-M’, respectively

This end stage of retinogenesis is the peak time point at which rods, bipolar cells and Müller glia are generated. To examine whether *Mettl3* deficiency affects retinal differentiation, we counted the numbers of rods, Müller glia and bipolar cells in the retina at p7, when the major retinogenesis phase was completed in the central retinas, even in *Mettl3^CKO^* mice. The results showed that the number of SOX9^+^ Müller glia was increased in *Mettl3^CKO^* retinas (Figure 2I, 2I’ and 2J). On the other hand, the number of VSX2^+^ bipolar cells remained relatively unchanged (Figure 2K, 2K’ and 2L), while the number of RHODOPSIN^+^ rods was reduced (Figure 2M, 2M’ and 2N). We also determined the cell densities of other retinal neurons. The numbers of BRN3A^+^ RGCs, CALBINDIN^+^ amacrine cells, and CAR^+^ cones were comparable between mutant and control retinas (Figure2-figure supplement 2), while the number of CALBINDIN^+^ horizontal cells was significantly reduced in the mutant retinas (Figure2-figure supplement 2E, 2E’ and 2G).

### Single-cell transcriptome analyses demonstrate distorted and extended late-stage retinogenesis in *Mettl3^CKO^* retinas

To examine how *Mettl3* deficiency affects gene expression in the retina, we performed RNA-seq analyses of control and *Mettl3^CKO^* retinas at p7, the time point at which the structural defects became obvious. In principal component analysis (PCA), mutant retinas were clearly separated from control retinas (Figure3-figure supplement 1A). Differentially expressed gene (DEG) analysis showed that 580 genes were upregulated and 532 genes were downregulated in *Mettl3^CKO^* retinas (Figure3-figure supplement 1B). Pearson analysis showed only subtle differences between control and mutant retinal transcriptomes (Figure3-figure supplement 1C), indicating that *Mettl3* delicately regulates retinal gene expression.

To examine the transcriptomic and cell status changes in *Mettl3^CKO^* retinal cells in a more comprehensive way, we performed single-cell RNA sequencing (scRNA-seq) on control and *Mettl3^CKO^* retinal cells at p7. A total of 5995 control and 6992 *Mettl3^CKO^* retinal cells passed quality control and were used for subsequent bioinformatic analysis. By applying unsupervised clustering and dimensional reduction and based on the expression of key marker genes, sequenced cells were clustered into nine clusters, representing the typical cellular composition of the retina at p7 (Figure 3A, 3B, and Figure3-figure supplement 2A). Rods, the largest cell population in the retina, occupied the largest cluster, and other retinal neurons, including cones, amacrine cells, bipolar cells, and Müller glial cells, each formed distinct clusters. Retinal ganglion cells and horizontal cells were not represented, probably due to their small population size. A group of cells along the trajectory toward bipolar cells and rods were judged as precursors for the two cell types based on their high expression of *Otx2* and *Neurod4*. A group of cells at one edge extending from Müller glia highly expressed cell cycle drivers, such as *Mki67*, *Pcna* and *Ccnb1*, suggesting that they were RPCs and were likely RPCs in the final phase of differentiation that reside on the retinal periphery at this time point; these cells were more abundantly observed in *Mettl3^CKO^* retinas whose retinogenesis period was extended (Figure3-figure supplement 3).

**Fig. 3.**
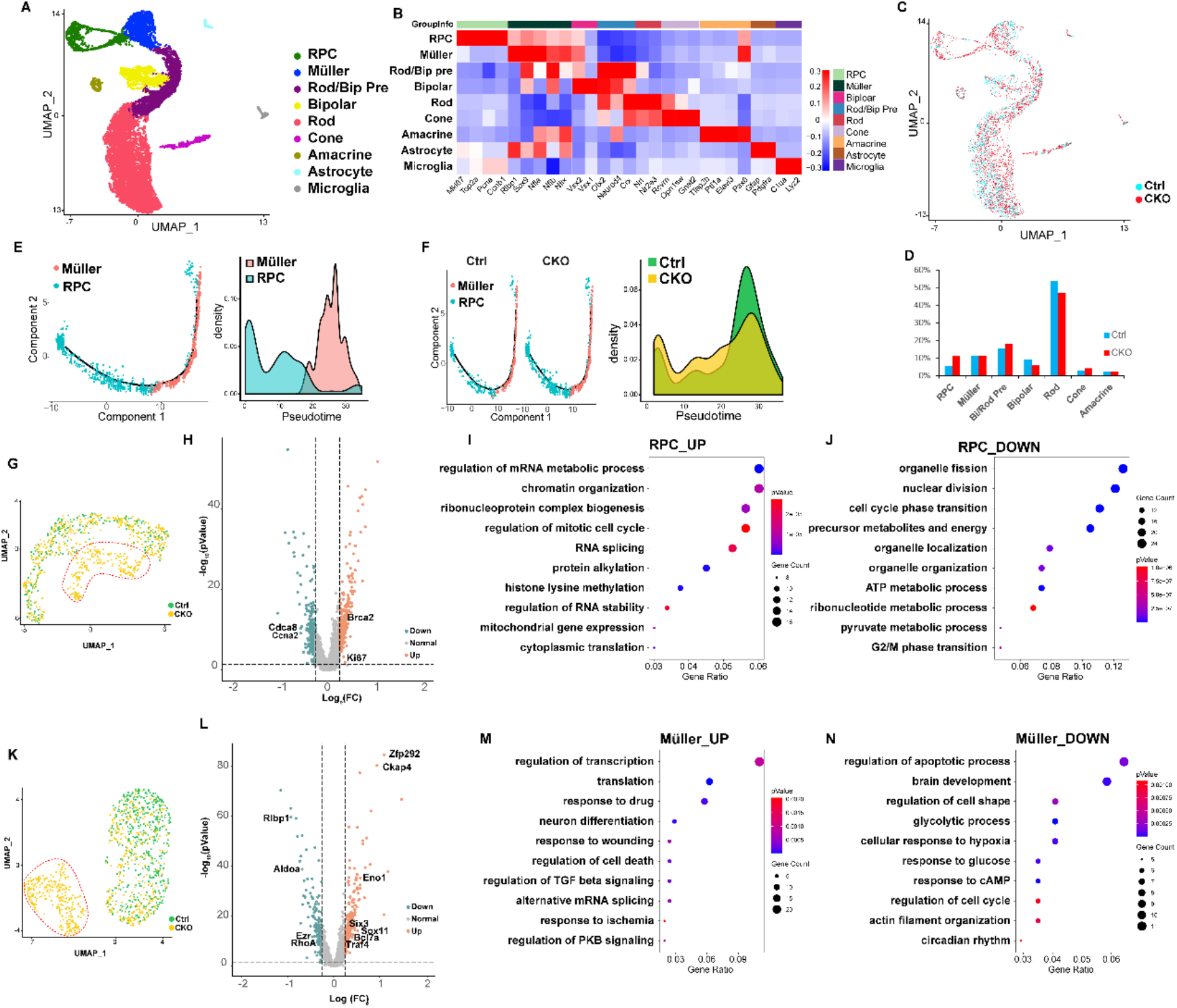
scRNA-seq analyses reveal distorted transcriptomes of late RPCs and Müller glial cells in *Mettl3^CKO^* retinas. (A) UMAP presenting the cell clusters of all sequenced cells. (B) Heatmap illustrating the expression patterns of key markers of each retinal cell type. (C) UMAP presenting the distribution of control and *Mettl3^CKO^* retinal cells. (D) Bar column presenting the cell compositions in control and *Mettl3^CKO^* retinas. (E) Left: pseudotime presentation illustrating the developmental progression from RPCs to Müller glia; right: cell density distributions of RPCs and Müller glia over the pseudotime period. (F) Left: separate pseudotime presentations of control and *Mettl3^CKO^* RPCs and Müller glia; right: cell density distributions of control and *Mettl3^CKO^* cells over the pseudotime period. (G) UMAP of reclustered RPCs. The red dotted line encircles a distinct cell cluster composed solely of *Mettl3^CKO^* RPCs. (H) Volcano plot showing gene expression differences between mutant RPCs (encircled by the red dotted line in A) and the remaining RPCs. (I) Bubble plot showing the biological processes enriched in genes that were upregulated in the *Mettl3*-mutant RPC cluster. (J) Bubble plot showing the biological processes enriched in genes that were downregulated in the *Mettl3*-mutant RPC cluster. (K) UMAP of reclustered Müller glia. The red dotted line encircles a distinct cell cluster composed solely of *Mettl3^CKO^* Müller glia. (L) Volcano plot showing gene expression differences between mutant Müller glia (encircled by the red dotted line in E) and the remaining Müller glia. (M) Bubble plot showing the biological processes enriched in genes that were upregulated in the *Mettl3*-mutant Müller glial cluster. (N) Bubble plot showing the biological processes enriched in genes that were downregulated in the *Mettl3*-mutant Müller glial cluster.

Control and *Mettl3^CKO^* retinal cells were largely mingled across clusters (Figure 3C). Cell number counting showed that the proportion of rods was slightly reduced in *Mettl3^CKO^* retinas and that the proportions of Müller glia in control and *Mettl3^CKO^* retinas were comparable, while the proportion of RPCs in *Mettl3^CKO^* retinas was nearly 2-fold higher than that in control retinas (Figure 3D and Figure3-figure supplement 2B), consistent with the extended retinogenesis observed for *Mettl3^CKO^* RPCs (Figure 2). Considering that RPCs at this age are near the end of retinogenesis, most would become Müller glia. As a consequence, the final proportion of Müller glia in *Mettl3^CKO^* retinas should be higher than that in control retinas, consistent with the manual counting of cells in the central retina at p7 (Figure 2). To observe the progression of this end-stage retinogenesis, we selected RPCs and Müller glia for pseudotime analysis. The results demonstrated a continuous cell status transition from RPCs to Müller glia (Figure 3E), consistent with the gliogenic status of RPCs at this time point. Analysis of the pseudotime distributions of control and *Mettl3^CKO^* cells revealed that the gliogenesis progression of *Mettl3^CKO^* RPCs lagged behind that of control RPCs (Figure 3F), suggesting delayed Müller gliogenesis in *Mettl3^CKO^* retinas, which is consistent with the extended retinogenesis in *Mettl3^CKO^* retinas revealed by BrdU labeling (Figure 2). Thus, the scRNA-seq examination of p7 control and *Mettl3^CKO^* retinal cells reveals distorted and extended late-stage retinogenesis in the mutant retina at the transcriptomic level.

### *Mettl3* deficiency distorts the transcriptomes of late RPCs and Müller glial cells

Clustering analysis of all the cells sequenced failed to reveal distinct cell clusters in the mutant retinas (Figure 3C). We suspected that subtle transcriptomic changes in some types of retinal cells were masked in the whole dataset. We thus performed reclustering of individual clusters of cells. The reclustering revealed distinct mutant clusters in the Müller, RPC, bipolar and rod populations (Figure 3G, 3K and Figure3-figure supplement 4A), especially for the Müller glia population, in which the mutant cell cluster was distantly separated from the control cell cluster (Figure 3K). Cell cycle phase analysis of RPCs showed that G2/M and S phase cells were distinctively distributed along two sides of UMAP, but the cell cycle phases were not responsible for the formation of the mutant cell cluster (Figure3-figure supplement 4B). Since phenotypic analyses indicated that RPCs and Müller glia were the two cell types most significantly affected by *Mettl3* deletion and that the mutant cell clusters were more pronounced in the RPC and Müller populations, we next focused on RPCs and Müller glia. DEG analysis of the mutant RPC cell cluster and the remaining RPC cells showed that 286 genes were upregulated and 212 genes were downregulated in mutant RPCs (Figure 3H). The upregulated genes were enriched for biological processes related to gene expression regulation, while downregulated genes were enriched for biological processes such as ‘organelle organization’ and ‘precursor metabolites and energy’ (Figure 3I and 3J). Importantly, key cell cycle machinery components, such as *Ki67* and *Brca2*, was upregulated, but many cell cycle regulators, such as *Ccna2* and *Cdca8*, were downregulated (Figure 3H-3J and Supplementary Table 1), which explained why the cell cycle progression of *Mettl3^CKO^* RPCs was distorted (Figure 2). DEG analysis of the mutant Müller cell cluster and the remaining Müller cells showed that 239 genes were upregulated and 181 genes were downregulated in mutant Müller cells (Figure 3L). The upregulated genes in the mutant Müller cell cluster were enriched for genes regulating transcription that have been linked with RPC development, such as *Six3* and *Sox11* (Figure 3L, 3M, and Supplementary Table 2). The downregulated genes in the mutant Müller cell cluster were enriched for regulators participating in ‘cell shape’ and ‘actin filament organization’, including important actin cytoskeleton regulators, such as *RhoA* and *Ezr* (Figure 3L, 3N, and Supplementary Table 2), indicating compromised physical properties of mutant Müller cells, which explained the distorted morphology of Müller cells and severely disorganized tissue structure in *Mettl3^CKO^* retinas (Figure 1). In addition, many genes that are essential for the supporting functions of Müller cells, such as those involved in the ‘glycolytic process’, including *Aldoa*, *Eno1*, and *Rlbp1*, were significantly downregulated in the mutant Müller cell cluster (Figure 3L, 3N, and Supplementary Table 2), indicating that the physiological functions of Müller glia in *Mettl3^CKO^* retinas were compromised, which likely contributes to the functional abnormality of *Mettl3^CKO^* retinas.

Taken together, reclustering and DEG analyses of scRNA-seq data reveal that *Mettl3* deficiency distorts the transcriptomes of late RPCs and Müller cells, which is likely the molecular basis for the distorted cellular behaviors of RPCs and Müller glia in *Mettl3^CKO^* retinas.

### The m^6^A epitranscriptome landscape of the mouse retina

To explore how m^6^A modification might be involved in retinal development and physiology, we next characterized the mouse retinal m^6^A epitranscriptome using MeRIP-seq. With a fold enrichment cutoff of 3, we detected 15471 peaks on 6735 genes, 13340 peaks on 6621 genes, and 14041 peaks on 5940 genes in p0, p6 and adult retinas, respectively (Figure 4A and Supplementary Table 3). The retinal m^6^A epitranscriptomes at different ages were similar to each other that of the 8421 m^6^A-modified genes detected in the retina, 53% were detected in all ages, and 76% were detected in at least two ages (Figure 4A). The retinal m^6^A peaks were largely localized in the CDS region, with pronounced enrichment around the stop codon (Figure 4B and Figure4-figure supplement 1A), and enriched for the ‘GGACU’ sequence (Figure 4C), consistent with the topological features observed in the m^6^A epitranscriptomes of other tissues (Dominissini et al., 2012; Meyer et al., 2012; Zaccara et al., 2019; Zhao et al., 2017). The mouse retinal m^6^A epitranscriptome was enriched for genes participating in biological processes related to retinal development, structural organization, and visual function, such as ‘synapse organization’, ‘axonogenesis’, ‘dendrite development’, ‘cell junction assembly’ and ‘retina development’, including many genes that have been demonstrated to play essential roles in retinal development, physiology or pathology (Figure 4D, 4E and Supplementary Table 4). The retinal m^6^A epitranscriptome at p0 was further enriched for genes involved in cell cycle progression, consistent with the proliferative status of the tissue at the time (Figure4-figure supplement 1B and Supplementary Table 4). Thus, the MeRIP-seq analyses revealed that the retinal m^6^A epitranscriptome is enriched for genes that play key roles in many essential activities of the retina and might participate in regulating these biological processes.

**Fig. 4.**
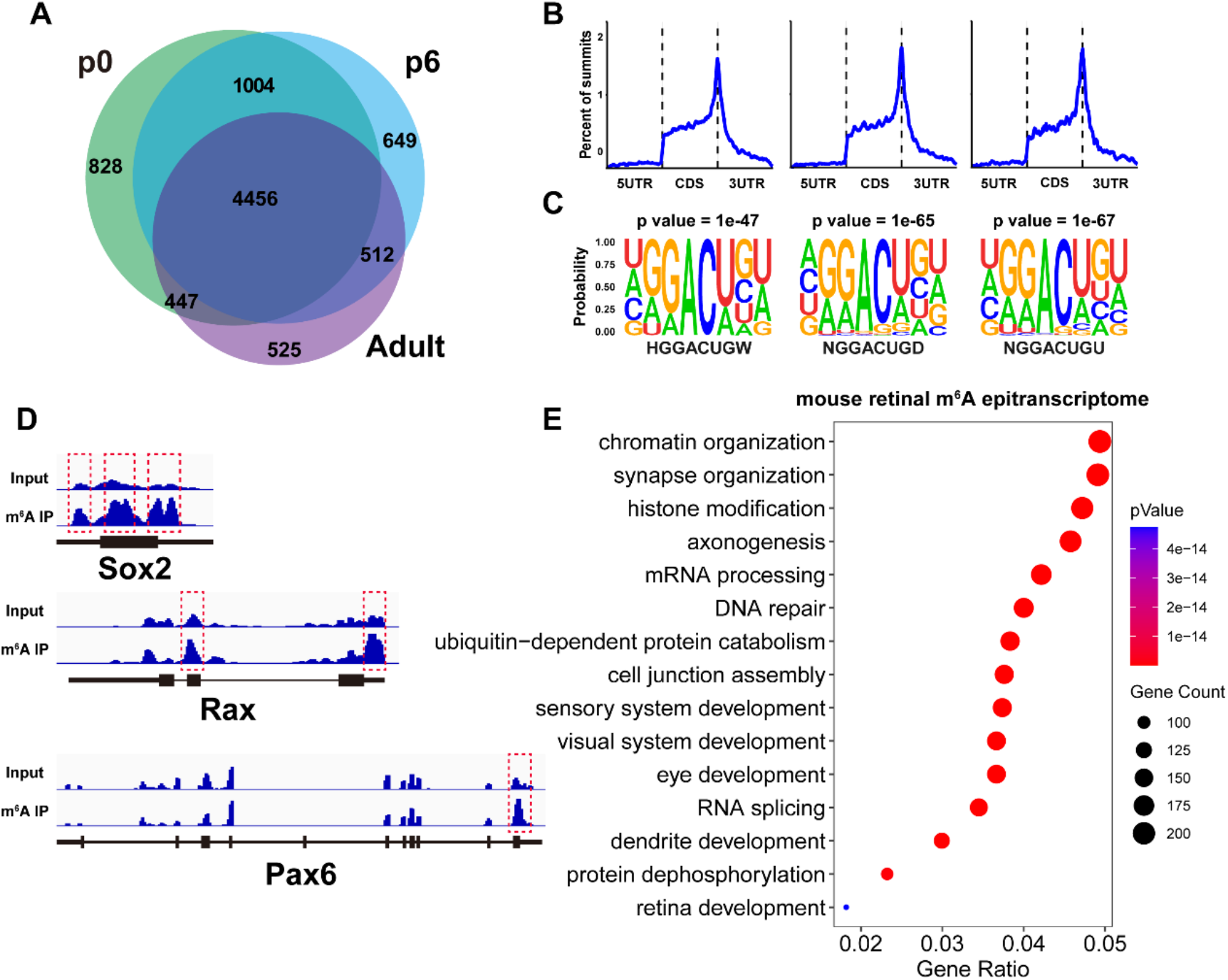
Mouse retinal m^6^A epitranscriptome. (A) Pie graph illustrating the number of genes carrying m^6^A modification detected by MeRIP-seq in the mouse retina at different ages. The numbers represent the number of genes in the respective pie region. (B) Plots illustrating the distribution of m^6^A peaks along mouse retinal transcripts. (C) The most enriched motif among m^6^A peak sequences in the mouse retina transcriptome. (D) IGV views illustrating the m^6^A peak distribution along transcripts of important retinal regulatory genes revealed by MeRIP-seq. (E) Bubble plot showing the biological processes enriched in genes carrying m^6^A modification detected in the retinas of mice at all ages.

### m^6^A modification promotes the degradation of RPC-enriched transcripts during late-stage retinogenesis

To investigate how m^6^A modification regulates the retinal transcriptome, we performed integrative analyses of the scRNA-seq data and the MeRIP-seq data. First, we analyzed the expression levels of m^6^A-tagged retinal transcripts in control and *Mettl3*-mutant cell clusters. Single-sample gene set enrichment analysis (ssGSEA) showed that m^6^A-tagged transcripts tended to be upregulated in *Mettl3*-mutant RPC and Müller glia clusters, and in *Mettl3*-mutant rod and bipolar cell clusters, but to a lesser extent (Figure 5A). Grouping genes based on the number of m^6^A peaks showed that as the number of m^6^A sites increased, the genes were upregulated to greater levels in *Mettl3*-mutant RPC and Müller glia clusters (Figure 5B). These results suggest that m^6^A modification promotes the degradation of modified transcripts in RPCs and Müller cells. Examining the m^6^A modification status of the DEG genes in *Mettl3*-mutant RPC and Müller glia clusters showed that most of the upregulated genes in *Mettl3*-mutant clusters were subject to m^6^A modification, while in contrast, a much smaller portion of the downregulated genes did (Figure 5C), suggesting that one major direct cause for gene upregulation in *Mettl3*-mutant RPC and Müller glia clusters was due to lack of m^6^A-mediated degradation.

**Fig. 5.**
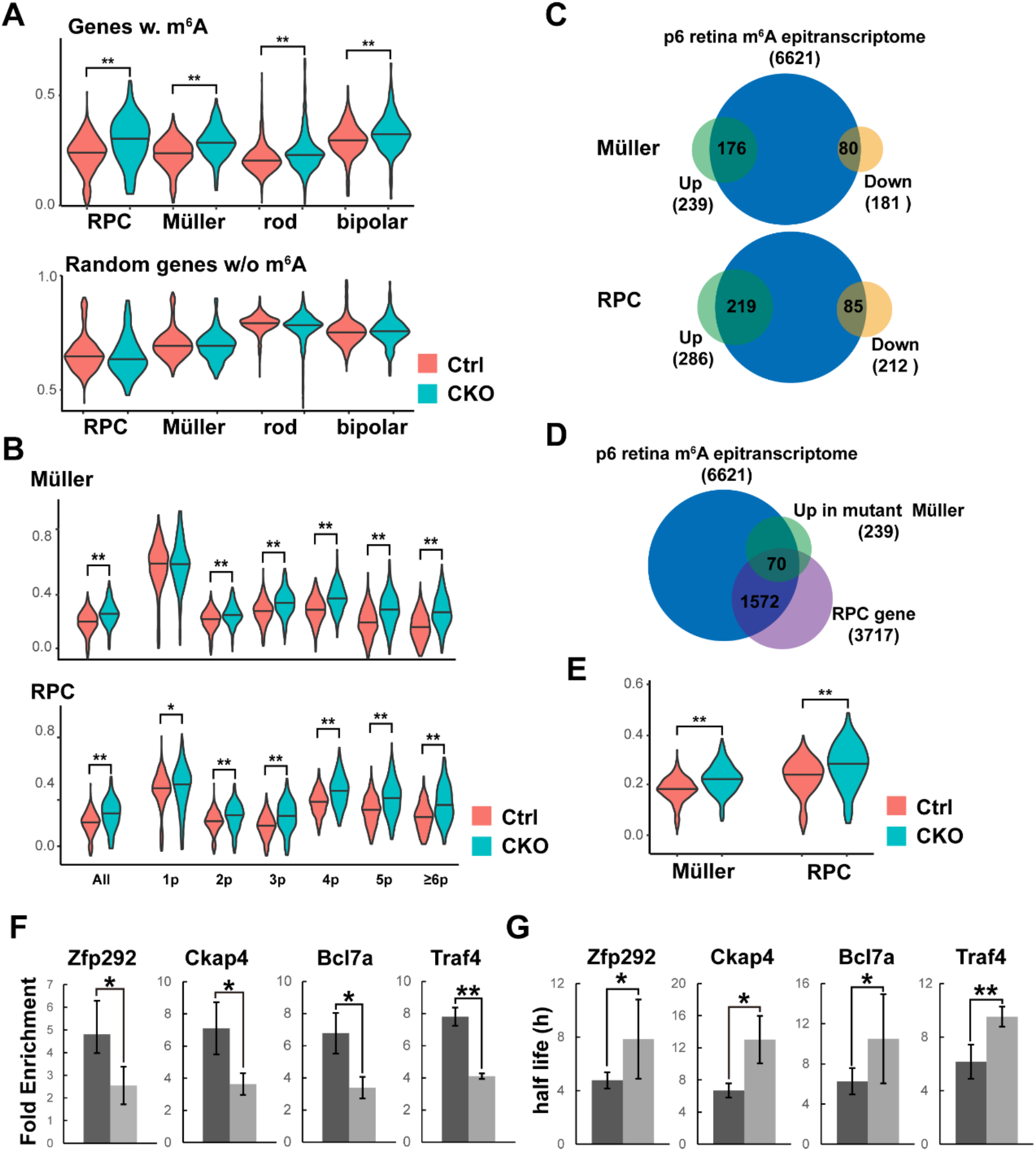
m^6^A modification promotes the degradation of RPC-enriched transcripts. (A) Top: Violin graph of ssGSEA scores illustrating the expression levels of m^6^A-modified genes in control and *Mettl3^CKO^* retinal cells. Bottom: Violin graph of ssGSEA scores of randomly selected genes that do not carry m^6^A to serve as controls. (B) Violin graphs of ssGSEA scores illustrating the expression levels of different groups of m^6^A-modified genes in control and *Mettl3^CKO^* Müller glia (top) and RPCs (bottom). The genes are grouped based on the number of m^6^A peaks along their transcript. (C) Pie graphs showing the number of m^6^A-modified genes that were upregulated (green pies) or downregulated (yellow pie) in *Mettl3*-mutant Müller glia (top) or RPC (bottom) clusters. The numbers inside the parentheses indicate the number of genes, and the numbers inside the figure indicate the number of overlapping genes between the two pies. (D) Pie graph showing the number of m^6^A-modified genes that were upregulated in the *Mettl3*-mutant Müller glial cluster (green pie) and enriched in the RPC transcriptome (purple pie). The numbers inside the parentheses indicate the number of genes, and the numbers inside the figure indicate the number of overlapping genes between the pies. (E) Violin graphs of ssGSEA scores illustrating the expression levels of m^6^A-modified RPC genes (purple pie in D) in the control and *Mettl3*-mutant Müller glia clusters and RPC clusters. (F) MeRIP-qPCR results revealed downregulation of m^6^A modification of the transcripts in *Mettl3^CKO^* retinas. (G) Transcript half-life measurements revealed prolonged half-lives of the transcripts in *Mettl3^CKO^* retinas. (H) * p < 0.05, ** p < 0.01.

As in most neural tissues, gliogenesis occurs toward the end of retinogenesis. By constitutively expressing a set of progenitor genes, Müller cells maintain a certain level of retinal regeneration potential (Blackshaw et al., 2004; Goldman, 2014; Jadhav et al., 2009). However, to function as a glial cell, Müller cells need to establish a unique transcriptome different from that of progenitors (Lin et al., 2019; Nelson et al., 2011). How this transcriptomic transition from progenitors to glial cells is achieved remains unexplored. We previously compared the transcriptomes of Müller glia and RPCs, yielding 3717 genes that are significantly more abundant in RPCs than in Müller glia(Lin et al., 2019). Among these 3717 RPC genes, 1642 were subject to m^6^A modification in the p6 retina. Among these 1642 m^6^A-tagged RPC-enriched genes, 70 were significantly upregulated in the *Mettl3*-mutant Müller cell cluster (Figure 5D and Supplementary Table 5). To systematically examine the expression levels of m^6^A-tagged RPC genes, we performed ssGSEA analyses of the 1642 m^6^A-tagged RPC genes between the *Mettl3*-mutant Müller cell cluster and the remaining Müller cells, which showed that these m^6^A-tagged RPC genes were significantly upregulated in the *Mettl3*-mutant Müller cell cluster (Figure 5E). In fact, these m^6^A-tagged RPC genes were already upregulated in the *Mettl3*-mutant late RPC cluster (Figure 5E). These results suggest that *Mettl3*-mutant Müller cells and RPCs maintain abnormal levels of the transcriptome of RPCs, which is likely due to the loss of the m^6^A modification in the transcripts.

To test whether m^6^A modification promotes the degradation of RPC transcripts, we selected 8 candidate genes (*Zfp292*, *Ckap4*, *Nxt1*, *Apex1*, *Bcl7a*, *Traf4*, *Rap2b*, and *Sulf2*) from the 70 m^6^A-tagged RPC-enriched genes that were significantly upregulated in the *Mettl3*-mutant Müller cluster (Figure 5D and Supplementary Table 5). All 8 genes carry multiple m^6^A tags around the stop codon (Figure5-figure supplement 1A and Supplementary Table 3) and are involved in various biological processes, such as transcription, cytoskeleton function, and nuclear export. Using MeRIP-qPCR, we confirmed that the m^6^A modification levels of the transcripts of these genes were reduced in *Mettl3^CKO^* retinas (Figure 5F and Figure5-figure supplement 1B). We next measured the half-lives of the transcripts of these genes in the control and *Mettl3^CKO^* retinas at p1. The results showed that the half-lives of the transcripts of five of these genes were extended significantly in *Mettl3^CKO^* retinas compared with those in the retinas of their littermate controls (Figure 5G and Figure5-figure supplement 1C), demonstrating that m^6^A modification promotes the degradation of these RPC transcripts in the retina.

Taken together, these analyses on the expression of m^6^A-tagged genes in control and *Mettl3*-mutant retinal cells demonstrate that m^6^A modification promotes the degradation of RPC-enriched transcripts during late-stage retinogenesis, which likely is essential for the timely termination of retinogenesis and the proper establishment of the transcriptome of Müller glial cells.

### Overexpression of m^6^A-tagged RPC genes in late RPCs disturbs late-stage retinogenesis

We next wanted to test how upregulation of m^6^A-tagged RPC genes would affect late-stage retinogenesis. For this purpose, we performed *in vivo* electroporation to overexpress the 5 candidate genes that showed extended transcript half-lives in *Mettl3^CKO^* retinas (Figure 5G and Figure5-figure supplement 1C) in p1 RPCs. At p3, two days after electroporation, we examined the cell cycle status of GFP-marked electroporated RPCs through PCNA staining. The results showed that there were significantly fewer PCNA^-^; GFP^+^ cells in the retinas electroporated with *Zfp292*-, *Ckap4*-, *Traf4*-, and *Bcl7a*-overexpression plasmids (Figure 6A and 6B), suggesting that overexpression of these four genes prohibited RPCs from exiting the cell cycle. We then traced the cell fates of GFP-marked electroporated RPCs at p14. The results showed that overexpression of *Zfp292*, *Ckap4*, *Traf4*, or *Bcl7a* led to an increase in the percentage of Müller cells at the expense of amacrine cells or rod photoreceptors (Figure 6C and 6D), suggesting that overexpression of these four genes biased the late RPCs to adopt Müller glial fate versus neuronal fates. The architecture of the retinas electroporated with *Zfp292*-, *Ckap4*-, *Traf4*-, and *Bcl7a*-overexpression plasmids was grossly normal, and phalloidin staining showed intact OLMs in these retinas (data not shown). We then coelectroporated all four genes in the retina. Phalloidin staining of the electroporated retinas at p14 showed that while the OLMs of these retinas were grossly intact, at a few regions, the OLMs are broken, and a few rods escaped to the subretinal space (Figure 6E, arrows in the insert image indicate the escaped rods). Thus, *in vivo* electroporation experiments demonstrated that ectopic overexpression of m^6^A-tagged RPC genes disturbs late-stage retinogenesis and structural homeostasis of the retina, reminiscent of the phenotypes observed in *Mettl3^CKO^* retinas, suggesting that proper downregulation of m^6^A-tagged RPC transcripts is essential for RPCs to exit retinogenesis and the establishment of the supporting functions of Müller cells, and thus the physiological homeostasis of the mature retina.

**Fig. 6.**
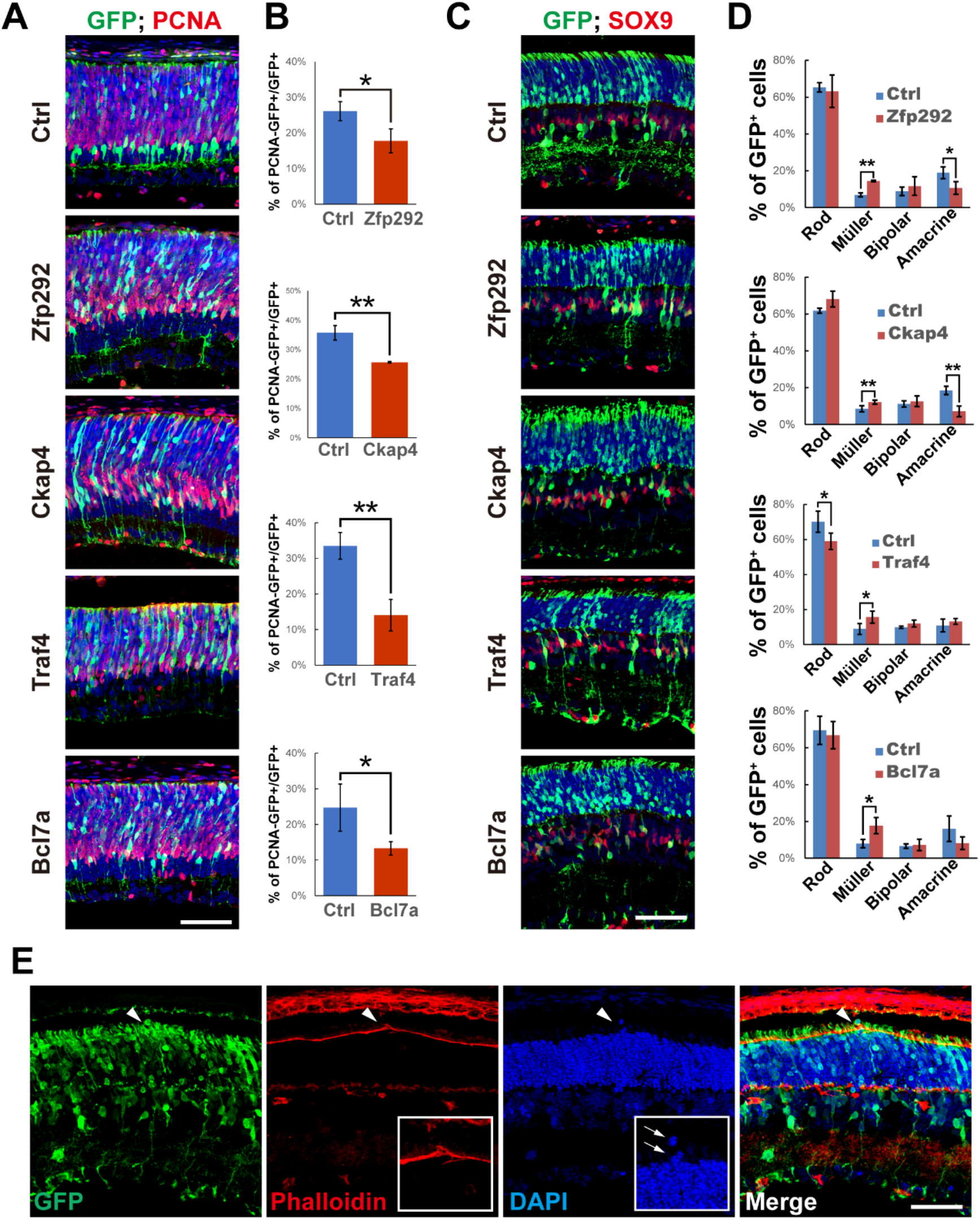
Overexpression of m^6^A-tagged RPC-enriched genes disturbs late-stage retinogenesis. (A) Confocal images of p3 retinas stained for GFP and PCNA. The retinas were transfected with plasmids overexpressing cDNAs of candidate genes via *in vivo* electroporation at p1. (B) Quantification of the percentage of GFP-labeled electroporated RPCs that had exited the cell cycle (PCNA^-^) 48 h after electroporation. (C) Confocal images of p14 retinas stained for GFP and SOX9. The retinas were transfected with plasmids overexpressing cDNAs of candidate genes via *in vivo* electroporation at p1. (D) Quantification of the cell compositions of GFP-labeled electroporated cells. (E) Confocal images of a retina stained for GFP and phalloidin. The retinas were transfected with a plasmid mix overexpressing cDNAs of *Zfp292*, *Ckap4*, *Traf4*, and *Bcl7a* via *in vivo* electroporation at p1 and were collected at p14. The arrowheads indicate a broken OLM region, which is further shown at higher magnification in the inserts. The arrows in the insert indicate escaped rods. The data in B and D are presented as the means ± standard deviations, corresponding to three independent biological replicates. * p < 0.05, ** p < 0.01. The scale bars in A, C, and E are 50 μm.

## Discussion

In this study, using *Mettl3^CKO^* mice as a model, we revealed that *Mettl3* is essential for late stage retinogenesis progression and structural and physiological homeostasis in the mature retina. Mechanistically, we showed that m^6^A promotes the degradation of RPC transcripts, thereby promoting the termination of retinogenesis and fine-tuning the transcriptomic transition from RPCs to Müller glial cells, the disruption of which leads to abnormalities in late stage retinogenesis and compromises the glial functions of Müller cells and thus the structural and physiological homeostasis of the mature retina.

One interesting finding of our study is that the regulatory function of *Mettl3* during retinal development is restricted to the late stage retinogenesis period. *Six3-Cre* mice express CRE recombinase in RPCs beginning from embryonic day 9.5 (Furuta et al., 2000); however, the retinas of *Mettl3^CKO^* mice remained normal until the postnatal period. Even though all are referred to as RPCs, RPCs continue to adjust their behavior during retinogenesis. For example, RPCs gradually go through a series of competent states to produce different types of retinal cells at different retinogenesis periods so that early RPCs produce retinal ganglion, cone, horizontal and some amacrine cells, while late RPCs produce rod, some amacrine, and bipolar cells before finally becoming Müller glial cells (Agathocleous and Harris, 2009; Cepko, 2014). Accompanying the gradual changes in the retinogenic behaviors, RPCs likely continue to adjust the gene regulatory networks to accommodate retinogenic tasks at different developmental stages. Indeed, scRNA-seq analyses of retinal development showed that the transcriptomes of early and late RPCs are distinct from each other (Clark et al., 2019). The postnatal retinogenesis period is the critical time when late RPCs quickly slow the cell cycle and finally exit the cell cycle to terminate retinogenesis in a timely fashion. Our study suggests that the m^6^A epitranscriptome delicately coordinates the transcriptomic adjustment needed for the timely termination of retinogenesis. In *Mettl3*-mutant RPCs, key cell cycle machinery components, such as *Ki67*, were upregulated, but many other cell cycle regulators were downregulated, which is likely the molecular basis for the distorted cell cycle. Furthermore, many m^6^A-tagged RPC transcripts were abnormally upregulated in *Mettl3*-mutant RPCs, which likely disrupts the balance of the gene regulatory networks in late RPCs. Indeed, overexpression of m^6^A-tagged RPC-enriched genes in late RPCs distorts the retinogenesis behavior of the cells. Thus, our study suggests that m^6^A delicately modulates the transcriptome of late RPCs to prepare them for the timely termination of retinogenesis.

In *Mettl3^CKO^* retinas, Müller gliogenesis was slightly enhanced, although delayed, which might be due to elevated RPC gene expression. However, the generated Müller cells were not functioning properly, which is also likely due to elevated RPC gene expression. As the last cell type generated by RPCs at the end of retinogenesis and as the major glial cell type in the retina, the close relationship between Müller glia and RPCs has long been noted. Many factors that play important roles in RPC proliferation and differentiation are also highly expressed in Müller glia, and several transcriptomic comparison studies have revealed similarities between Müller glia and RPCs at the transcriptome level (Blackshaw et al., 2004; Nelson et al., 2011; Roesch et al., 2008). Encouraged by these cues, the regenerative potential of Müller glia has been intensively explored in recent years (Goldman, 2014; Hoang et al., 2020; Jorstad et al., 2017; Lahne et al., 2020; Yao et al., 2018; Zhou et al., 2020). However, to function as glial cells, Müller cells must develop a unique transcriptome that differs from that of RPCs to accommodate their structural and physiological supporting roles in the adult neural retina (Lin et al., 2019; Nelson et al., 2011), but how this delicate transcriptome transition is achieved is unclear. In *Mettl3^CKO^* retinas, mutant Müller cells abnormally upregulated a group of RPC transcripts. Our results suggest that loss of m^6^A-mediated transcript degradation is a major cause for the upregulation of RPC transcripts in *Mettl3*-mutant Müller cells. Abnormal upregulation of RPC transcripts likely collectively disturbs the gene regulatory networks in Müller cells, leading to compromised glial function of Müller cells and thus abnormalities in the structure and visual function of the mature retina. Thus, our study demonstrates an interesting epitranscriptomic mechanism fine-tuning the transcriptomic transition from RPCs to Müller glial cells.

As a derivative of the neural tube, the neural retina shares similar regulatory principles with other central neural tissues. Similar to Müller cells in the retina, the supporting functions and regeneration potentials of glial cells in the central nervous system are hotly debated neural biological questions (Allen and Lyons, 2018; Guo et al., 2014; Qian et al., 2020; Wang et al., 2021; Zuchero and Barres, 2015). It will be interesting to test whether m^6^A plays similar roles during the development of glial cells in other nervous systems. It is also worth future studies to determine whether m^6^A is involved in the cell fate reprogramming between glial cells and neurons.

## Materials and Methods

### Animals

All animal studies were performed in accordance with the protocol approved by the Institutional Animal Care and Use Committee of Zhongshan Ophthalmic Center, and all animals were housed in the animal care facility of Zhongshan Ophthalmic Center. *Six3-Cre* mice were kindly provided by Dr. Furuta from the University of Texas(Furuta et al., 2000). *Mettl3^floxed^* mice were kindly provided by Dr. Tong Minghan from Shanghai Institute of Biochemistry and Cell Biology, Chinese Academy of Sciences(Lin et al., 2017). *Rlbp1-GFP* mice were kindly provided by Edward M. Levine from the University of Utah(Vazquez-Chona et al., 2009).

### Histology, immunostaining and images

For cryosectioning, eyes were fixed overnight in 4% formaldehyde, cryopreserved with 15% and then 30% sucrose, frozen in Tissue-Tek OCT freezing medium and sectioned using a cryostat microtome (Leica CM1950). For paraffin sectioning, eyes were fixed with Davidson’s Fixative overnight, processed, embedded in paraffin, and sectioned using a microtome (Leica RM2235). Before subsequent histological staining, paraffin sections underwent a standard procedure for deparaffinization, and cryosections were air-dried for 5 mins and then soaked in PBS for 5 min to wash off OCT. H&E staining was performed using a standard procedure. For fluorescent immunostaining, section slides were first heated in citrate buffer at 95°C for 30 mins for antigen retrieval. After cooling to room temperature, slides were incubated with primary antibodies prepared in PBST with 5% fetal bovine serum at 4°C overnight, washed with PBST, incubated with fluorophore-conjugated secondary antibodies, washed, counterstained with 4’,6-diamidino-2-phenylindole (DAPI), and finally mounted with cover glasses using VECTASHIELD mounting medium (Vector Labs). The following antibodies were used: rabbit anti-METTL3 (abCam, ab195352, RRID: AB_2721254), rabbit anti-RECOVERIN (Millipore, AB5585, RRID: AB_2253622), sheep anti-VSX2 (Millipore, AB9014, RRID: AB_262173), rabbit anti-SOX9 (Millipore, AB5535, RRID: AB_2239761), rabbit anti-CALBINDIN (Swant, CB38), HPC-1 (mouse anti-SYNTAXIN, S0664, RRID: AB_477483) (Sigma), mouse anti-BRN3A (Millipore, MAB1585, RRID: AB_94166), chicken anti-GFP (abCam, ab13970, RRID: AB_300798), rabbit anti-N-CADHERIN (Santa Cruz, sc-7939, RRID: AB_647794), mouse anti-GS (BD Transduction Laboratories, AB_397880, RRID: AB_397880), mouse anti-GFAP (Sigma, G3893, RRID: AB_477010), mouse anti-BrdU (BD Biosciences, 347580, RRID: AB_10015219), rabbit anti-KI67 (abCam, ab15580, RRID: AB_443209), mouse anti-pH3 (Ser10) (Cell Signaling Technology, 9706, RRID: AB_331748), Alexa Fluor 488 (568) Donkey anti-mouse (ThermoFisher), Alexa Fluor 488 (568) Donkey anti-rabbit (ThermoFisher), Alexa Fluor 488 Donkey anti-chicken (Jackson ImmunoResearch), Alexa Fluor 488 (568) Donkey anti-sheep (ThermoFisher). Bright field H&E-stained images were taken under an inverted microscope (Zeiss Axio Observer). Immunofluorescence images were taken under a confocal microscope (Zeiss LSM 880 or LSM 980).

### BrdU labeling

Pups were injected subcutaneously with BrdU (0.32 μmol/g body weight). Two hours, or 6 h, or 48 h later, pups were sacrificed, and eyes were collected, fixed with 4% formaldehyde overnight, and subjected to cryosectioning and immunofluorescent staining as described above. After confocal images were taken, the number of BrdU^+^ cells was counted for each selected 200 μm retinal region.

### mfERG

The experiments were performed on p14 control (n=4) and *Mettl3^CKO^* (n=2) mice. The mice were anesthetized using sodium pentobarbital and positioned on a warming table to maintain body temperature. For each animal, only the right eye was examined. Before recordings started, the eyes were dilated with 0.5% tropicamide (SANTEN pharmaceutical, Japan) and local narcotized with 0.5% proparacaine hydrochloride (Ruinian Best, China). The eyes were positioned 1-2 mm in front of the device (Roland RETImap SLO, Roland, Germany). The animal Goldring recording electrode was placed at the corneal limbus. Subcutaneous silver needle electrodes served as reference and ground electrodes. An optical correction lens was positioned in front of the recording electrode. Viscous 2% methocel gel was applied between the cornea and contact lens. The stimuli were generated by a projector with a refresh rate of 60 Hz. The number of recording hexagons was set at 19, and the optic nerve head (ONH) was adjusted near the edge of the recording area. The stimulation parameter of light was set at mfERG Max. Twelve cycles were averaged for a final result.

### RNA-seq

RNA-seq was performed on 4 control and 4 CKO retinal samples. The retinas of p6 control and CKO pups were collected directly into TRIzol (ThermoFisher), and total RNA was extracted following the manufacturer’s instructions. Sequencing libraries were generated using the NEBNext Ultra RNA Library Prep Kit for Illumina (NEB) following the manufacturer’s instructions. The libraries were sequenced on an Illumina HiSeq platform, and 150 bp paired-end reads were generated. Raw data were filtered to remove low-quality reads, and the clean reads were mapped to the mouse genome (GRCm38, mm10) using Hisat2 v2.0.5. featureCounts v1.5.0-p3 was used to count the read numbers mapped to each gene, and then the FPKM of each gene was calculated. Differential expression analysis was performed using the DESeq2 R package. The resulting P values were adjusted using Benjamini and Hochberg’s approach. Genes with an adjusted P value < 0.05 were considered differentially expressed.

### scRNA-seq

Two retinas from 2 control pups and 2 retinas from 2 *Mettl3^CKO^* pups of the same litter were used for the scRNA-seq experiment. Retinas were digested with Papain (Worthington) to single cells and resuspended in PBS containing 0.04% BSA, and 2 retinal samples for each experimental group were mixed together as one sample for subsequent scRNA-seq. The scRNA-seq was performed by NovelBio Co., Ltd. Libraries were generated using the 10X Genomics Chromium Controller Instrument and Chromium Single Cell 3’ V3.1 Reagent Kits (10X Genomics). Libraries were sequenced using an Illumina sequencer (Illumina) on a 150 bp paired-end run.

We applied fastp with default parameter filtering of the adaptor sequence and removed the low-quality reads to obtain clean data. Then, the feature-barcode matrices were obtained by aligning reads to the mouse genome (GRCm38, mm10) using CellRanger v3.1.0. Cells containing over 200 expressed genes and a mitochondrial UMI rate below 20% passed the quality filtering, and mitochondrial genes were removed from the expression table. The Seurat package v3.1 was used for cell normalization and regression to obtain the scaled data. PCA was constructed based on the scaled data with the top 2000 highly variable genes, and the top 10 principal components were used for UMAP construction. Utilizing the graph-based cluster method, we acquired unsupervised cell clusters. Judged by the key markers expressed by each cluster, we merged some clusters and annotated the final clusters.

For pseudotime analysis, we applied single-cell trajectory analysis using Monocle2 using DDR-Tree and default parameters. To identify differentially expressed genes among samples, the function FindMarkers with the Wilcox rank sum test algorithm was used under the following criteria: lnFC > 0.25, P value < 0.05, min.pct > 0.1. To characterize the relative activity of gene sets with different m^6^A features, we performed QuSAGE (2.16.1) analysis. Fisher’s exact test was applied to identify the significant GO terms and FDR was used to correct the p-values.

### MeRIP-seq

The retinas of C57BL/6J mice were collected directly into TRIzol (Thermo Fisher). Total RNA was extracted according to the manufacturer’s instructions, and RNAs from 14 retinas were pooled together for each MeRIP-seq. MeRIP-seq was performed according to a published protocol(Dominissini et al., 2013). Briefly, PolyA-mRNA was purified using a Dynabeads mRNA DIRECT Purification kit (Thermo Fisher). mRNA was fragmented in fragmentation buffer heated at 94°C for 5 mins. Fragmented mRNA was immunoprecipitated against an m^6^A antibody (Synaptic Systems) bound to Protein G Dynabeads (Thermo Fisher) at 4°C for 2 h, washed, and eluted by competition with free m^6^A (Sigma). After recovery of precipitated m^6^A-mRNA, sequencing libraries were constructed using the VAHTS Universal V6 RNA-seq Library Prep Kit (Vazyme). The libraries were sequenced on an Illumina HiSeq platform on a 150 bp paired-end run. Raw data were filtered to remove low-quality reads to obtain clean reads. Clean reads were mapped to the mouse genome (GRCm38, mm10) using BWA mem v0.7.12. m^6^A peaks were determined using MACS2 v2.1.0 with the q-value threshold of enrichment set at 0.05. Motif analysis was done using MEME. GO term analysis was performed using clusterProfiler package (Yu et al., 2012).

### MeRIP-qPCR

Total RNA from retinas (5-6 retinas per sample) was extracted using TRIzol. MeRIP for qPCR was performed as described above for MeRIP-seq except that the mRNA fragmentation step was skipped. m^6^A pull-down and 1% reserved input RNAs were reversed transcribed into cDNAs using the SuperScript II First-strand Synthesis System for RT-PCR (Thermo Fisher). qPCR was performed using the SYBR green-based method on a LightCycler 480 (Roche). The primers were listed in the Supplementary table 6. Enrichment of genes in m^6^A-tagged genes in pull-down samples was calculated based on ΔCt between the pull-down sample and the input sample, and was normalized to an internal negative control, *Rsp14*, which does not carry m^6^A modification.

### Molecular cloning

Full-length *Zfp292*, *Ckap4*, *Traf4*, *Bcl7a*, *Rap2b*, *Nxt1*, *Apex*, and *Sulf2* cDNAs were PCR amplified from a cDNA pool derived from embryonic mouse retina total RNA and cloned into the pCAGIG vector (gift from Dr. Connie Cepko and obtained through Addgene).

### Mouse retina *in vivo* electroporation

Newborn mouse pups were anesthetized by chilling on ice. A small incision was made in the eyelid and sclera near the edge of the cornea with a 30-gauge needle (Hamilton). Purified plasmid solutions (2-5 μg/μl) in PBS were injected into the subretinal space through the previous incision using a Hamilton syringe with a 32-gauge blunt-ended needle under a dissecting microscope. After DNA injection, a tweezer-type electrode was placed to softly hold the pup heads, and five 80 V pulses of 50 ms duration and 950 ms intervals were applied using a Gemini X2 pulse generator (BTX). After electroporation, pups were allowed to recover on a warming pad until they regained consciousness and were returned to their mother. The pups were sacrificed at p14, and the retinas were collected and subjected to cryosection as described above. Eye sections were immunostained for GFP (to recognize transduced cells and their progenies) and SOX9 (to recognize Müller cells). For cell composition analysis of the electroporated cells, GFP^+^ cells in the ONL were judged as rods, GFP^+^; SOX9^+^ cells were judged as Müller cells, the GFP^+^ cells in the INL above the SOX9+ Müller cells (close to the OPL layer) were judged as bipolar cells, while the GFP^+^ cells in the INL below the SOX9^+^ Müller cells (close to the IPL layer) were judged as amacrine cells.

### RNA half-life measurement

*Mettl3^CKO^* and control mice were sacrificed at p1, and the retinas were collected and soaked in DMEM supplemented with actinomycin D (5 μm, Sigma) to inhibit transcription and cultured in a CO2 incubator at 37°C. The retinas were collected directly into TRIzoL at 0 h, 4 h, 8 h, and 16 h after drug treatment. Total RNA was extracted according to the manufacturer’s instructions. Two micrograms of total RNA from each sample was reverse transcribed into cDNA using SuperScript II (Thermo Fisher) and subjected to qPCR measurements using SYBR Green mix (Roche) and LightCycler LC480 (Roche). The primers were listed in table 1. The relative mRNA concentrations at each time point against time point 0 were calculated, the ln^mRNA^ ^concentration^ at time points 0, 4, 8, and 16 h were plotted to perform a linear regression analysis as a function of time, and the slope of the line was identified as the decay rate (k). The half-life was calculated with the following formula: t (1/2) = ln2/k(Chen et al., 2008).

### Statistical analysis

All experiments subjected to statistical analysis were performed with at least three biological replicates. Quantification data are presented as the mean ± standard deviation and the significance of differences between samples was assessed using Student’s t test.

### Data availability

The accession numbers for the RNA-seq, scRNA-seq and MeRIP-seq data are NCBI Gene Expression Omnibus: GSE180815, GSE181095, and GSE188607.

## Acknowledgments

We thank Dr. Minghan Tong, from Shanghai Institute of Biochemistry and Cell Biology, Chinese Academy of Sciences, for providing the *Mettl3^floxed^* mice, Dr. Edward M. Levine, from the University of Utah, for providing the *Rlbp1-GFP* mice, Dr. Yasuhide Furata, from the University of Texas, for providing the *Six3-Cre* mice. We thank the staff of Laboratory Animal Center at State Key Laboratory of Ophthalmology, Zhongshan Ophthalmic Center for technical support. This study was supported by the National Natural Science Foundation of China (81870659 and 81721003).

## Supplementary Figure legends

**Figure1- figure supplement 1.**
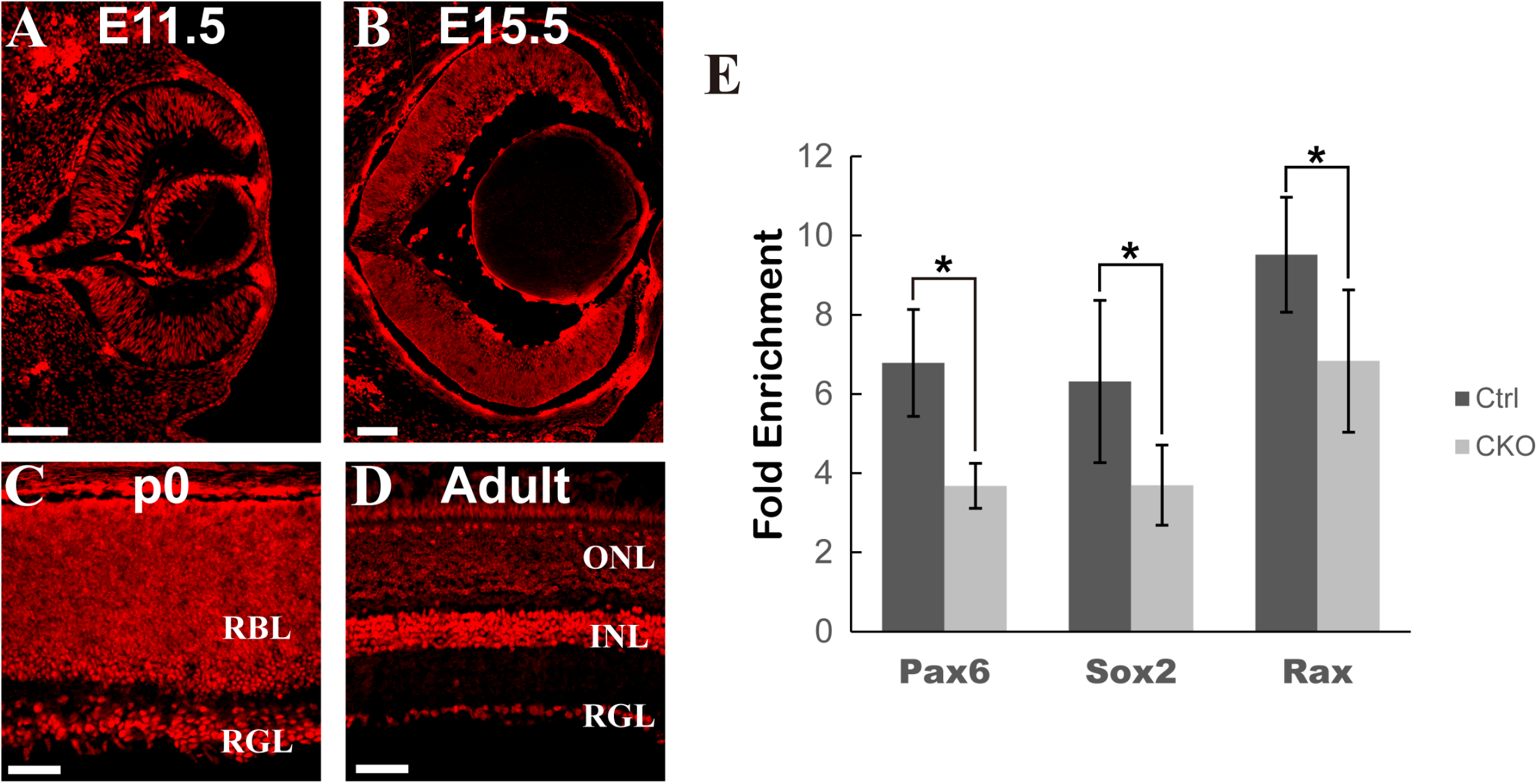
The expression of *Mettl3* in the mouse retina. (A-D) Confocal images of METTL3 expression in mouse retinas at different developmental stages. The scale bars are 100 μm. (E) MeRIP-qPCR results showing that the m^6^A levels of several candidate genes were significantly reduced in *Mettl3^CKO^* retinas. The data are presented as the means ± standard deviations, corresponding to three independent biological replicates. * p < 0.05.

**Figure1- figure supplement 2.**
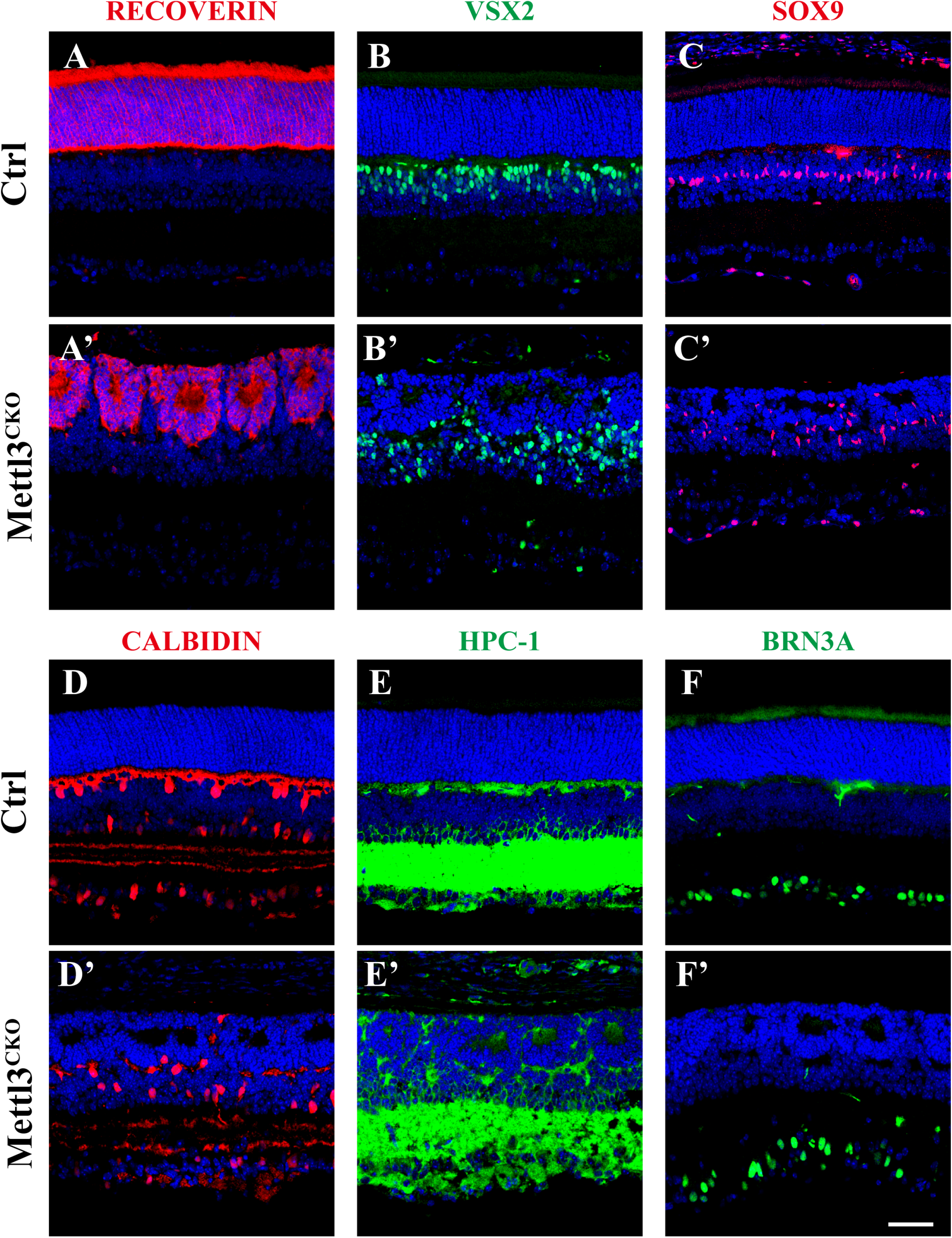
All types of retinal cells are generated in *Mettl3^CKO^* retinas. Confocal images of p14 retinas stained for RECOVERIN to illustrate photoreceptor cells (A, A’), VSX2 to illustrate bipolar cells (B, B’), SOX9 to illustrate Müller cells (C, C’), CALBINDIN to illustrate horizontal cells and subtype amacrine cells (D, D’), HPC-1 to illustrate amacrine cells (E, E’), and BRN3A to illustrate RGCs (F, F’). The scale bar in F’ is 50 μm and applies to all images in the figure.

**Figure2- figure supplement 1.**
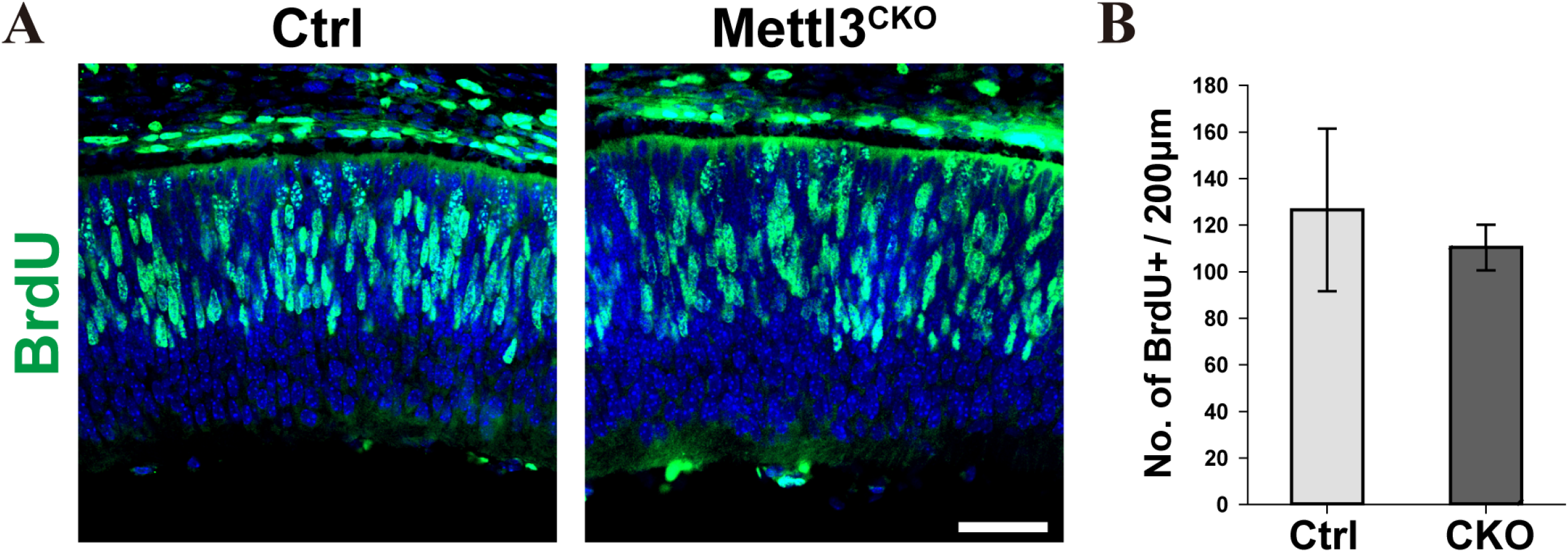
*Mettl3* deficiency does not affect the proliferation of early RPCs. (A) Confocal images of control and *Mettl3^CKO^* retinas at E15.5 stained for BrdU. The mothers were injected with BrdU, and the embryos were collected 2 h later. (B) Quantification of the BrdU^+^ cells in A and A’. The data are represented as the means ± standard deviations, corresponding to three independent biological replicates.

**Figure2- figure supplement 2.**
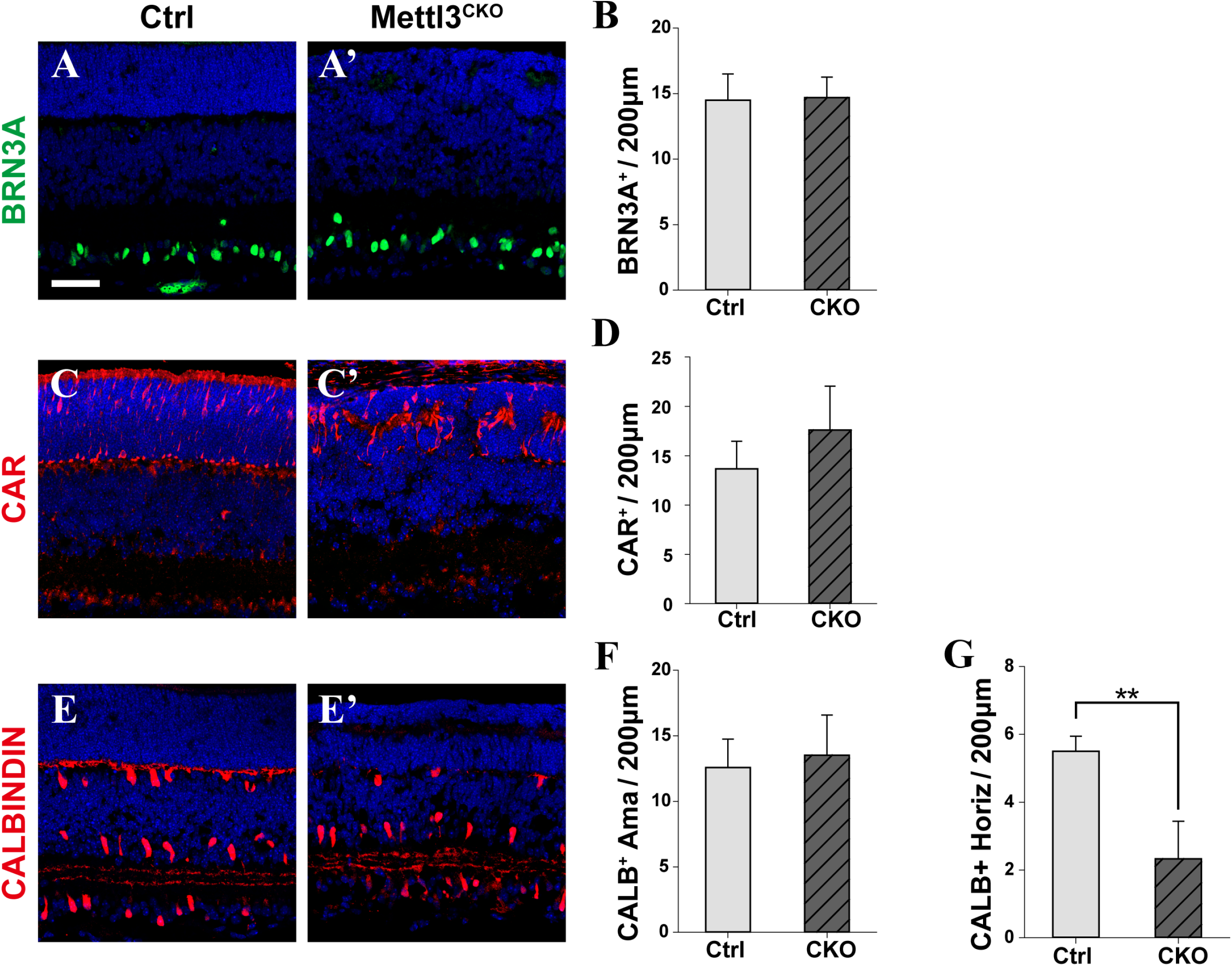
Different types of retinal cells at p7. (A, A’) Confocal images of p7 retinas stained for BRN3A to illustrate RGCs. (B) Quantification of the BRN3A^+^ cells in A and A’. (C, C’) Confocal images of p7 retinas stained for CAR to illustrate cones. (D) Quantification of the CAR^+^ cells in C and C’. (E, E’) Confocal images of p7 retinas stained for CALBINDIN to illustrate amacrine cells and horizontal cells. (F) Quantification of the CALBINDIN^+^ amacrine cells (cells close to the IPL) in E and E’. (G) Quantification of the CALBINDIN^+^ horizontal cells (cells close to the OPL) in E and E’. The data in B, D, F, and G are represented as the means ± standard deviations, corresponding to three independent biological replicates. ** p < 0.01. The scale bar in A is 50 μm and applies to all images in the figure.

**Figure3- figure supplement 1.**
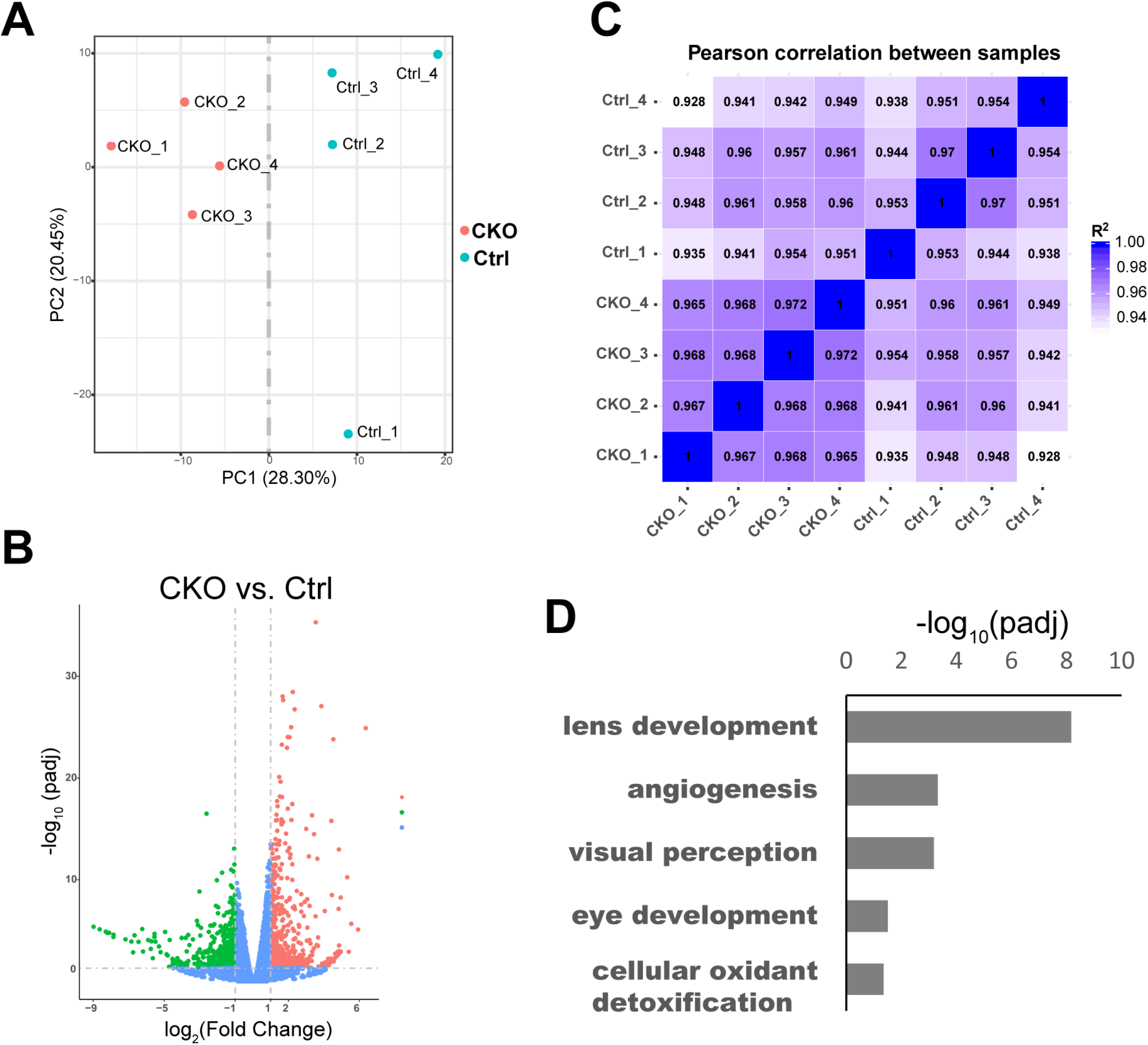
RNA-seq analyses of the transcriptomes of control and *Mettl3^CKO^* retinas. (A) PCA map illustrating separation of the control and *Mettl3^CKO^* retinal transcriptomes. (B) Volcano plot illustrating the gene expression differences between control and *Mettl3^CKO^* retinas. (C) Pearson scores of the transcriptomes of different samples. (D) Bar graph showing the biological processes enriched in the gene group that were downregulated in *Mettl3^CKO^* retinas.

**Figure3- figure supplement 2.**
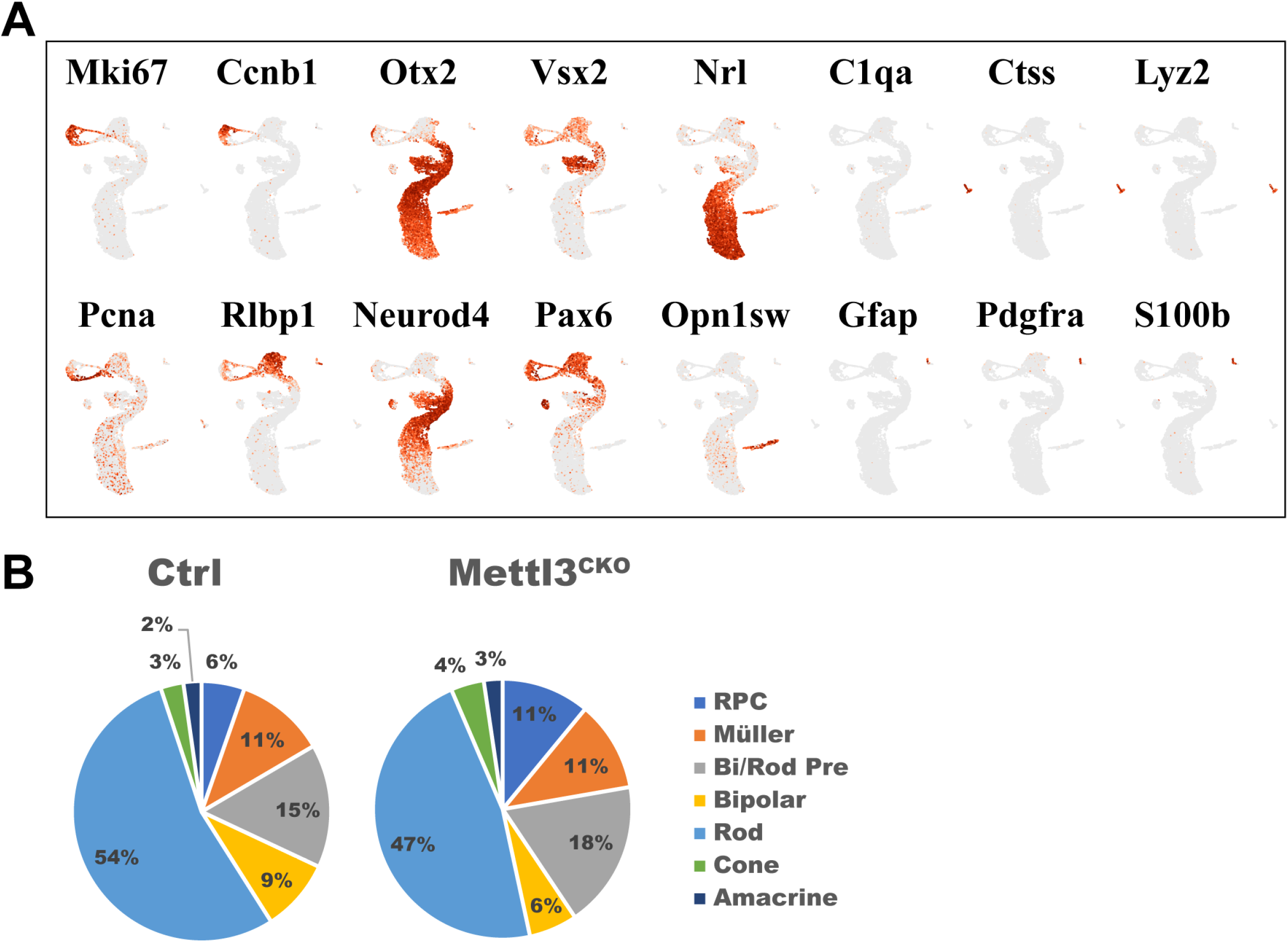
scRNA-seq analysis of *Mettl3^CKO^* retinal cells. (A) Expression patterns of marker genes of various types of retinal cells in UMAP. (B) Pie charts illustrating the cell type compositions in control and *Mettl3^CKO^* retinas.

**Figure3- figure supplement 3.**
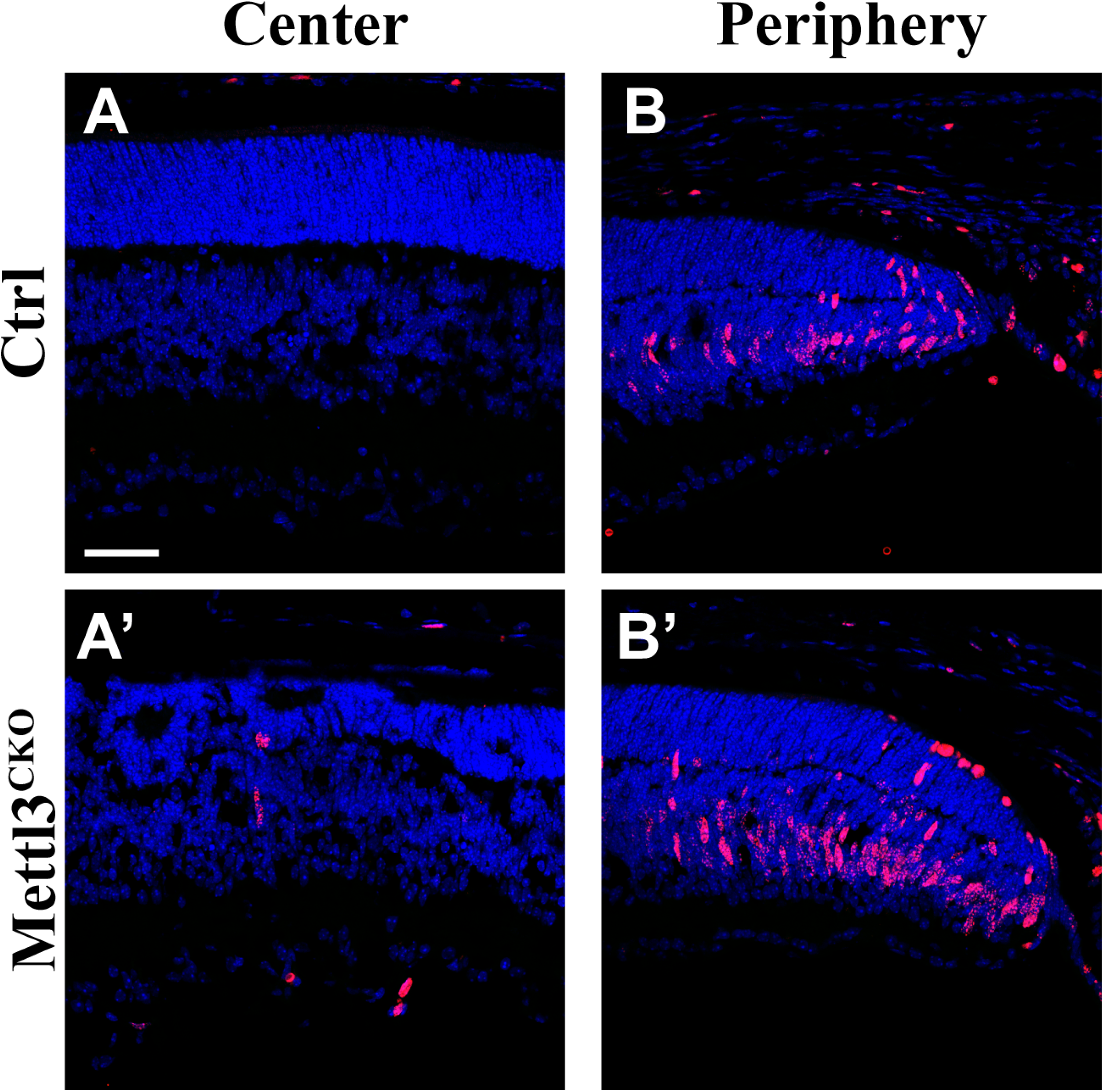
Proliferating RPCs in the retinas at p7. Confocal images of p7 control and *Mettl3^CKO^* retinas stained for KI67 to illustrate proliferating RPCs. A and A’ are central regions of the retinas, while B and B’ are peripheral regions of the retinas. The scale bar in A is 50 μm and applies to all images in the figure.

**Figure3- figure supplement 4.**
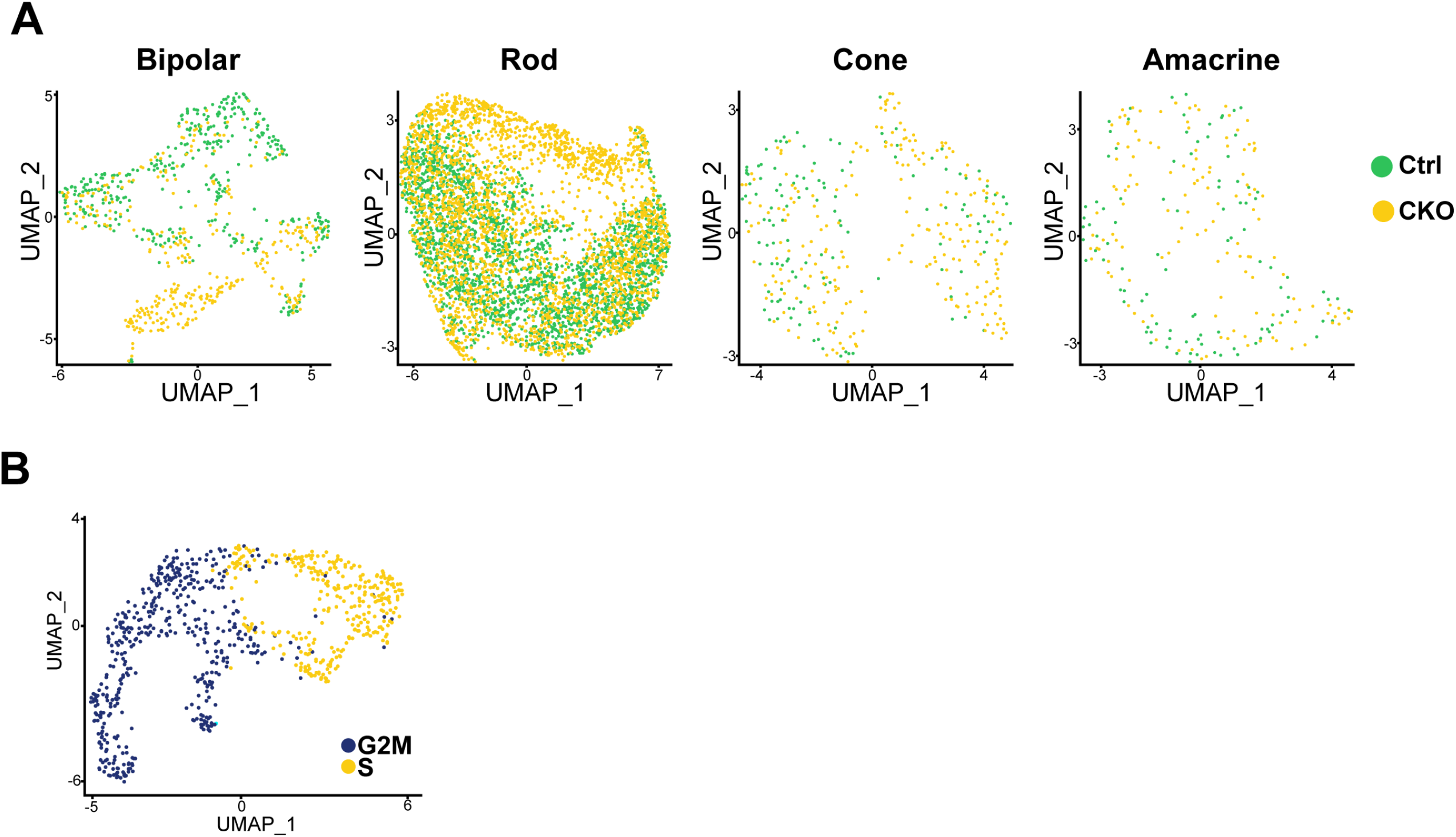
Reclustering of different types of retinal cells. (A) UMAPs of the reclustered bipolar cell, rod, cone and amacrine cell populations. (B) cell cycle phase distribution pattern in reclustered RPCs.

**Figure4- figure supplement 1.**
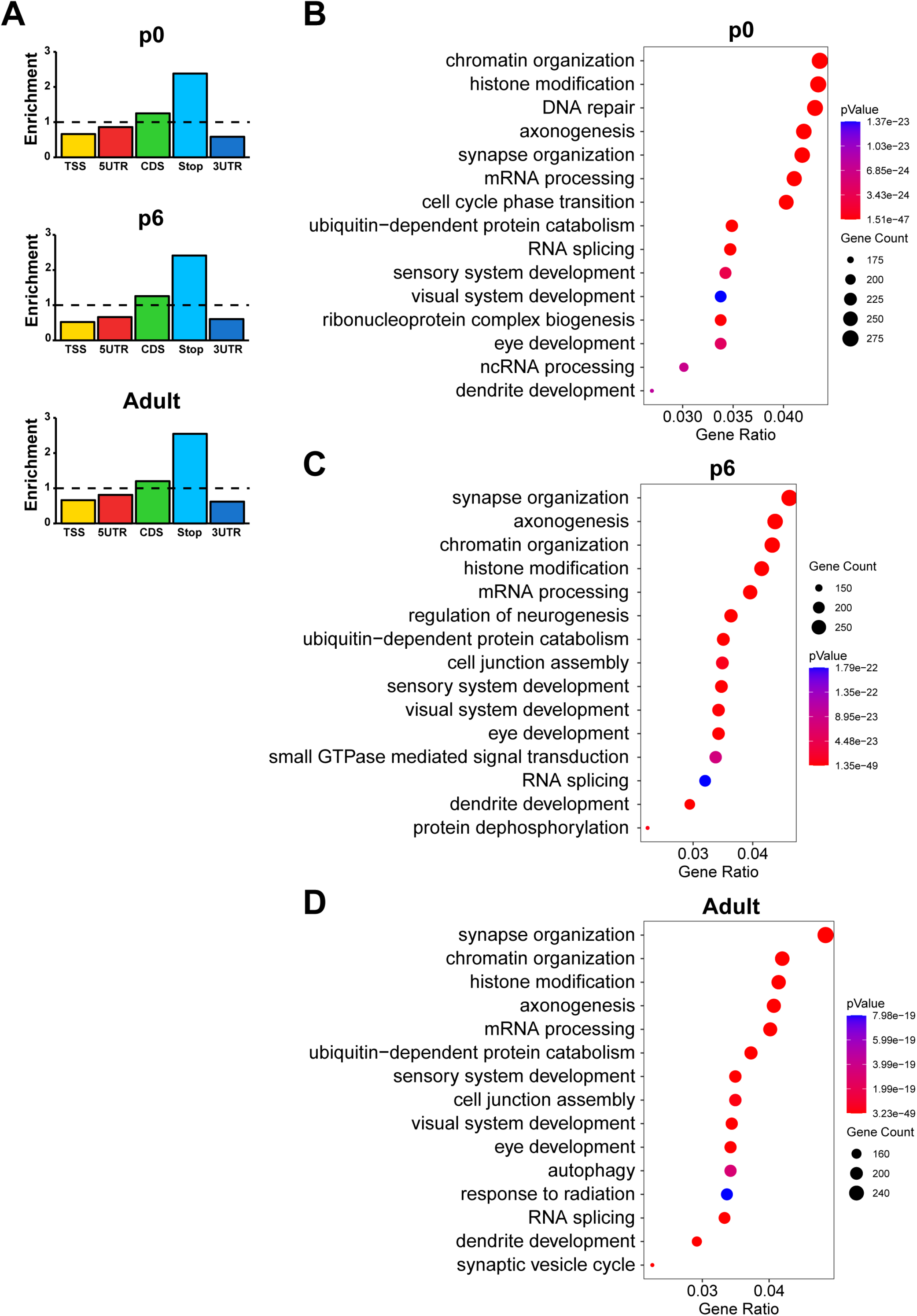
m^6^A epitranscriptomes of the mouse retina at different ages. (A) Bar graphs illustrating the enrichment status of m^6^A peaks in different regions of mouse retinal transcripts. (B-D) Bubble plots showing the biological processes enriched in genes carrying m^6^A modification detected in the retinas at p0 (B), p6 (C), and in adults (D).

**Figure5- figure supplement 1.**
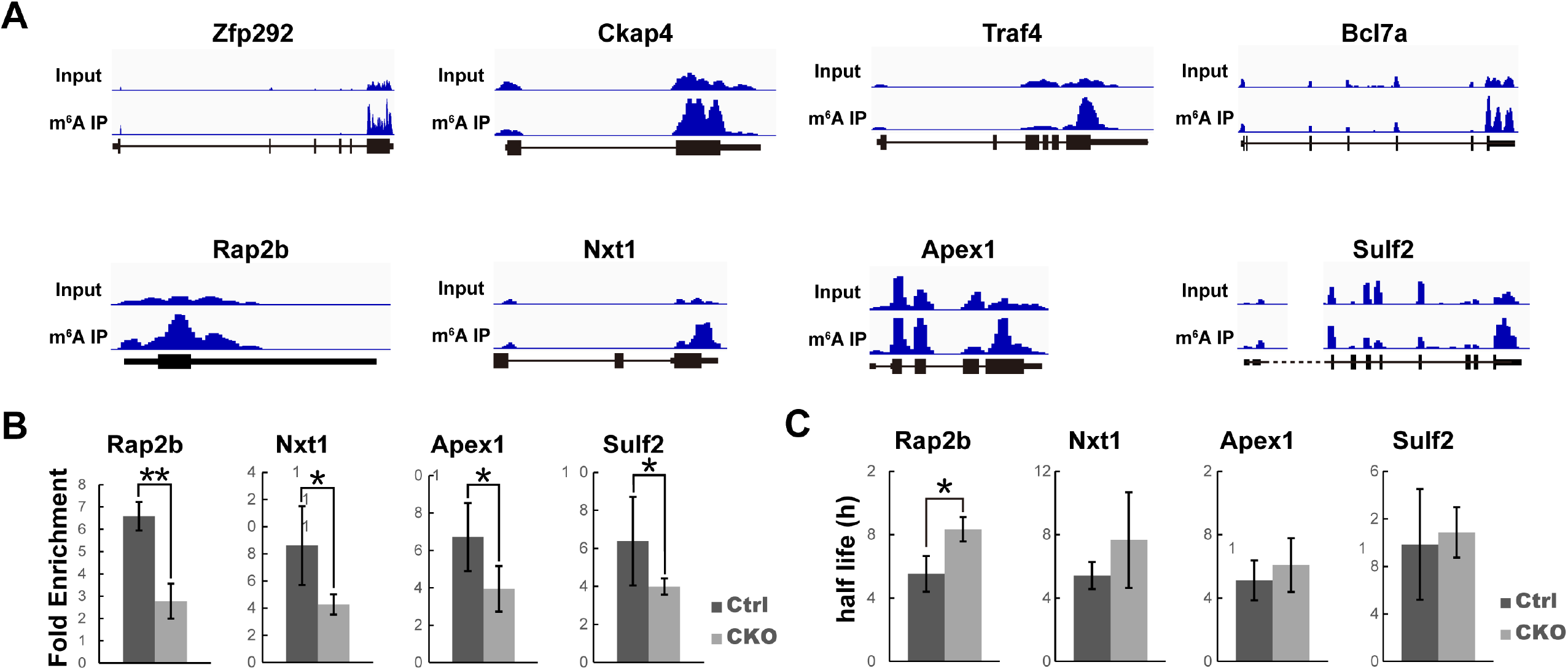
m^6^A-modified RPC-enriched genes. (A) IGV views illustrating the m^6^A peak distribution along transcripts of RPC-enriched genes as revealed by MeRIP-seq. (B) MeRIP-qPCR results revealing downregulation of m^6^A modification of the transcripts in *Mettl3^CKO^* retinas. (C) Transcript half-life measurements of the RPC-enriched genes carrying m^6^A modification. The data are presented as the means ± standard deviations, corresponding to three independent biological replicates. * p < 0.05, ** p < 0.01.

## Supplementary Tables for this manuscript

Supplementary Table 1. List of DEG genes in the *Mettl3*-mutant RPC cluster

Supplementary Table 2. List of DEG genes in the *Mettl3*-mutant Müller cell cluster

Supplementary Table 3. Mouse retinal m^6^A epitranscriptome

Supplementary Table 4. Biological processes enriched in the mouse retinal m^6^A epitranscriptome

Supplementary Table 5. List of RPC-enriched genes carrying m^6^A modification and upregulated in the *Mettl3*-mutant Müller cell cluster

Supplementary Table 6. List of primers

## References

Agathocleous, M., and Harris, W.A. (2009). From progenitors to differentiated cells in the vertebrate retina. Annual review of cell and developmental biology 25, 45–69. https://doi.org/10.1146/annurev.cellbio.042308.113259

Allen, N.J., and Lyons, D.A. (2018). Glia as architects of central nervous system formation and function. Science 362, 181–185. https://doi.org/10.1126/science.aat0473

Bassett, E.A., and Wallace, V.A. (2012). Cell fate determination in the vertebrate retina. Trends in neurosciences 35, 565–573. https://doi.org/10.1016/j.tins.2012.05.004

Blackshaw, S., Harpavat, S., Trimarchi, J., Cai, L., Huang, H., Kuo, W.P., Weber, G., Lee, K., Fraioli, R.E., Cho, S.H., Yung, R., Asch, E., Ohno-Machado, L., Wong, W.H., and Cepko, C.L. (2004). Genomic analysis of mouse retinal development. PLoS biology 2, E247.

Bokar, J.A., Rath-Shambaugh, M.E., Ludwiczak, R., Narayan, P., and Rottman, F. (1994). Characterization and partial purification of mRNA N6-adenosine methyltransferase from HeLa cell nuclei. Internal mRNA methylation requires a multisubunit complex. The Journal of biological chemistry 269, 17697–17704. https://doi.org/S0021-9258(17)32497-3

Bokar, J.A., Shambaugh, M.E., Polayes, D., Matera, A.G., and Rottman, F.M. (1997). Purification and cDNA cloning of the AdoMet-binding subunit of the human mRNA (N6-adenosine)-methyltransferase. RNA 3, 1233–1247.

Byrne, L.C., Khalid, F., Lee, T., Zin, E.A., Greenberg, K.P., Visel, M., Schaffer, D.V., and Flannery, J.G. (2013). AAV-mediated, optogenetic ablation of Muller Glia leads to structural and functional changes in the mouse retina. PloS one 8, e76075. https://doi.org/10.1371/journal.pone.0076075

Cepko, C. (2014). Intrinsically different retinal progenitor cells produce specific types of progeny. Nat Rev Neurosci 15, 615–627. https://doi.org/10.1038/nrn3767

Chen, C.Y., Ezzeddine, N., and Shyu, A.B. (2008). Messenger RNA half-life measurements in mammalian cells. Methods in enzymology 448, 335–357. https://doi.org/10.1016/S0076-6879(08)02617-7

Clark, B.S., Stein-O’Brien, G.L., Shiau, F., Cannon, G.H., Davis-Marcisak, E., Sherman, T., Santiago, C.P., Hoang, T.V., Rajaii, F., James-Esposito, R.E., Gronostajski, R.M., Fertig, E.J., Goff, L.A., and Blackshaw, S. (2019). Single-Cell RNA-Seq Analysis of Retinal Development Identifies NFI Factors as Regulating Mitotic Exit and Late-Born Cell Specification. Neuron 102, 1111–1126 e1115. https://doi.org/10.1016/j.neuron.2019.04.010

Dominissini, D., Moshitch-Moshkovitz, S., Salmon-Divon, M., Amariglio, N., and Rechavi, G. (2013). Transcriptome-wide mapping of N(6)-methyladenosine by m(6)A-seq based on immunocapturing and massively parallel sequencing. Nature protocols 8, 176–189. https://doi.org/10.1038/nprot.2012.148

Dominissini, D., Moshitch-Moshkovitz, S., Schwartz, S., Salmon-Divon, M., Ungar, L., Osenberg, S., Cesarkas, K., Jacob-Hirsch, J., Amariglio, N., Kupiec, M., Sorek, R., and Rechavi, G. (2012). Topology of the human and mouse m6A RNA methylomes revealed by m6A-seq. Nature 485, 201–206. https://doi.org/10.1038/nature11112

Frye, M., and Blanco, S. (2016). Post-transcriptional modifications in development and stem cells. Development (Cambridge, England) 143, 3871–3881. https://doi.org/10.1242/dev.136556

Furuta, Y., Lagutin, O., Hogan, B.L., and Oliver, G.C. (2000). Retina- and ventral forebrain-specific Cre recombinase activity in transgenic mice. Genesis 26, 130–132.

Goldman, D. (2014). Muller glial cell reprogramming and retina regeneration. Nat Rev Neurosci 15, 431–442. https://doi.org/10.1038/nrn3723

Guo, Z., Zhang, L., Wu, Z., Chen, Y., Wang, F., and Chen, G. (2014). In vivo direct reprogramming of reactive glial cells into functional neurons after brain injury and in an Alzheimer’s disease model. Cell Stem Cell 14, 188–202. https://doi.org/10.1016/j.stem.2013.12.001

Heavner, W., and Pevny, L. (2012). Eye development and retinogenesis. Cold Spring Harbor perspectives in biology 4. https://doi.org/10.1101/cshperspect.a008391

Hoang, T., Wang, J., Boyd, P., Wang, F., Santiago, C., Jiang, L., Yoo, S., Lahne, M., Todd, L.J., Jia, M., Saez, C., Keuthan, C., Palazzo, I., Squires, N., Campbell, W.A., Rajaii, F., Parayil, T., Trinh, V., Kim, D.W., Wang, G., Campbell, L.J., Ash, J., Fischer, A.J., Hyde, D.R., Qian, J., and Blackshaw, S. (2020). Gene regulatory networks controlling vertebrate retinal regeneration. Science. https://doi.org/10.1126/science.abb8598

Jadhav, A.P., Roesch, K., and Cepko, C.L. (2009). Development and neurogenic potential of Muller glial cells in the vertebrate retina. Prog Retin Eye Res 28, 249–262. https://doi.org/10.1016/j.preteyeres.2009.05.002

Jia, G., Fu, Y., Zhao, X., Dai, Q., Zheng, G., Yang, Y., Yi, C., Lindahl, T., Pan, T., Yang, Y.G., and He, C. (2011). N6-methyladenosine in nuclear RNA is a major substrate of the obesity-associated FTO. Nat Chem Biol 7, 885–887. https://doi.org/10.1038/nchembio.687

Jorstad, N.L., Wilken, M.S., Grimes, W.N., Wohl, S.G., VandenBosch, L.S., Yoshimatsu, T., Wong, R.O., Rieke, F., and Reh, T.A. (2017). Stimulation of functional neuronal regeneration from Muller glia in adult mice. Nature 548, 103–107. https://doi.org/10.1038/nature23283

Lahne, M., Nagashima, M., Hyde, D.R., and Hitchcock, P.F. (2020). Reprogramming Muller Glia to Regenerate Retinal Neurons. Annu Rev Vis Sci 6, 171–193. https://doi.org/10.1146/annurev-vision-121219-081808

Lee, Y., Choe, J., Park, O.H., and Kim, Y.K. (2020). Molecular Mechanisms Driving mRNA Degradation by m(6)A Modification. Trends Genet 36, 177–188. https://doi.org/S0168-9525(19)30268-9

Lin, S., Guo, J., and Chen, S. (2019). Transcriptome and DNA Methylome Signatures Associated With Retinal Muller Glia Development, Injury Response, and Aging. Invest Ophthalmol Vis Sci 60, 4436–4450. https://doi.org/10.1167/iovs.19-27361

Lin, Z., Hsu, P.J., Xing, X., Fang, J., Lu, Z., Zou, Q., Zhang, K.J., Zhang, X., Zhou, Y., Zhang, T., Zhang, Y., Song, W., Jia, G., Yang, X., He, C., and Tong, M.H. (2017). Mettl3-/Mettl14-mediated mRNA N(6)-methyladenosine modulates murine spermatogenesis. Cell Res 27, 1216–1230. https://doi.org/10.1038/cr.2017.117

Liu, J., Harada, B.T., and He, C. (2019). Regulation of Gene Expression by N(6)-methyladenosine in Cancer. Trends Cell Biol 29, 487–499. https://doi.org/10.1016/j.tcb.2019.02.008

Liu, J., Yue, Y., Han, D., Wang, X., Fu, Y., Zhang, L., Jia, G., Yu, M., Lu, Z., Deng, X., Dai, Q., Chen, W., and He, C. (2014). A METTL3-METTL14 complex mediates mammalian nuclear RNA N6-adenosine methylation. Nat Chem Biol 10, 93–95. https://doi.org/10.1038/nchembio.1432

Livneh, I., Moshitch-Moshkovitz, S., Amariglio, N., Rechavi, G., and Dominissini, D. (2020). The m(6)A epitranscriptome: transcriptome plasticity in brain development and function. Nat Rev Neurosci 21, 36–51. https://doi.org/10.1038/s41583-019-0244-z

MacDonald, R.B., Randlett, O., Oswald, J., Yoshimatsu, T., Franze, K., and Harris, W.A. (2015). Muller glia provide essential tensile strength to the developing retina. The Journal of cell biology 210, 1075–1083. https://doi.org/10.1083/jcb.201503115

Masland, R.H. (2001). The fundamental plan of the retina. Nature neuroscience 4, 877–886. https://doi.org/10.1038/nn0901-877

Meyer, K.D., and Jaffrey, S.R. (2014). The dynamic epitranscriptome: N6-methyladenosine and gene expression control. Nature reviews 15, 313–326. https://doi.org/10.1038/nrm3785

Meyer, K.D., and Jaffrey, S.R. (2017). Rethinking m(6)A Readers, Writers, and Erasers. Annual review of cell and developmental biology 33, 319–342. https://doi.org/10.1146/annurev-cellbio-100616-060758

Meyer, K.D., Saletore, Y., Zumbo, P., Elemento, O., Mason, C.E., and Jaffrey, S.R. (2012). Comprehensive analysis of mRNA methylation reveals enrichment in 3’ UTRs and near stop codons. Cell 149, 1635–1646. https://doi.org/10.1016/j.cell.2012.05.003

Nelson, B.R., Ueki, Y., Reardon, S., Karl, M.O., Georgi, S., Hartman, B.H., Lamba, D.A., and Reh, T.A. (2011). Genome-wide analysis of Muller glial differentiation reveals a requirement for Notch signaling in postmitotic cells to maintain the glial fate. PloS one 6, e22817. https://doi.org/10.1371/journal.pone.0022817

Newman, E., and Reichenbach, A. (1996). The Muller cell: a functional element of the retina. Trends in neurosciences 19, 307–312. https://doi.org/0166-2236(96)10040-0 [pii]

Qian, H., Kang, X., Hu, J., Zhang, D., Liang, Z., Meng, F., Zhang, X., Xue, Y., Maimon, R., Dowdy, S.F., Devaraj, N.K., Zhou, Z., Mobley, W.C., Cleveland, D.W., and Fu, X.D. (2020). Reversing a model of Parkinson’s disease with in situ converted nigral neurons. Nature 582, 550–556. https://doi.org/10.1038/s41586-020-2388-4

Roesch, K., Jadhav, A.P., Trimarchi, J.M., Stadler, M.B., Roska, B., Sun, B.B., and Cepko, C.L. (2008). The transcriptome of retinal Muller glial cells. J Comp Neurol 509, 225–238. https://doi.org/10.1002/cne.21730

Roundtree, I.A., Evans, M.E., Pan, T., and He, C. (2017). Dynamic RNA Modifications in Gene Expression Regulation. Cell 169, 1187–1200. https://doi.org/S0092-8674(17)30638-4

Shen, W., Fruttiger, M., Zhu, L., Chung, S.H., Barnett, N.L., Kirk, J.K., Lee, S., Coorey, N.J., Killingsworth, M., Sherman, L.S., and Gillies, M.C. (2012). Conditional Mullercell ablation causes independent neuronal and vascular pathologies in a novel transgenic model. J Neurosci 32, 15715–15727. https://doi.org/10.1523/JNEUROSCI.2841-12.2012

Shi, H., Wei, J., and He, C. (2019). Where, When, and How: Context-Dependent Functions of RNA Methylation Writers, Readers, and Erasers. Molecular cell 74, 640–650. https://doi.org/S1097-2765(19)30317-X

Sledz, P., and Jinek, M. (2016). Structural insights into the molecular mechanism of the m(6)A writer complex. Elife 5. https://doi.org/10.7554/eLife.18434

Vazquez-Chona, F.R., Clark, A.M., and Levine, E.M. (2009). Rlbp1 promoter drives robust Muller glial GFP expression in transgenic mice. Invest Ophth Vis Sci 50, 3996–4003. https://doi.org/10.1167/iovs.08-3189

Vecino, E., Rodriguez, F.D., Ruzafa, N., Pereiro, X., and Sharma, S.C. (2016). Glia-neuron interactions in the mammalian retina. Prog Retin Eye Res 51, 1–40. https://doi.org/10.1016/j.preteyeres.2015.06.003

Wang, J., O’Sullivan, M.L., Mukherjee, D., Punal, V.M., Farsiu, S., and Kay, J.N. (2017). Anatomy and spatial organization of Muller glia in mouse retina. J Comp Neurol 525, 1759–1777. https://doi.org/10.1002/cne.24153

Wang, L.L., Serrano, C., Zhong, X., Ma, S., Zou, Y., and Zhang, C.L. (2021). Revisiting astrocyte to neuron conversion with lineage tracing in vivo. Cell 184, 5465–5481 e5416. https://doi.org/S0092-8674(21)01052-7

Wang, P., Doxtader, K.A., and Nam, Y. (2016a). Structural Basis for Cooperative Function of Mettl3 and Mettl14 Methyltransferases. Molecular cell 63, 306–317. https://doi.org/10.1016/j.molcel.2016.05.041

Wang, X., Feng, J., Xue, Y., Guan, Z., Zhang, D., Liu, Z., Gong, Z., Wang, Q., Huang, J., Tang, C., Zou, T., and Yin, P. (2016b). Structural basis of N(6)-adenosine methylation by the METTL3-METTL14 complex. Nature 534, 575–578. https://doi.org/10.1038/nature18298

Yao, K., Qiu, S., Wang, Y.V., Park, S.J.H., Mohns, E.J., Mehta, B., Liu, X., Chang, B., Zenisek, D., Crair, M.C., Demb, J.B., and Chen, B. (2018). Restoration of vision after de novo genesis of rod photoreceptors in mammalian retinas. Nature 560, 484–488. https://doi.org/10.1038/s41586-018-0425-3

Yu, G., Wang, L.G., Han, Y., and He, Q.Y. (2012). clusterProfiler: an R package for comparing biological themes among gene clusters. OMICS 16, 284–287. https://doi.org/10.1089/omi.2011.0118

Zaccara, S., Ries, R.J., and Jaffrey, S.R. (2019). Reading, writing and erasing mRNA methylation. Nature reviews 20, 608–624. https://doi.org/10.1038/s41580-019-0168-5

Zhao, B.S., Roundtree, I.A., and He, C. (2017). Post-transcriptional gene regulation by mRNA modifications. Nature reviews 18, 31–42. https://doi.org/10.1038/nrm.2016.132

Zheng, G., Dahl, J.A., Niu, Y., Fedorcsak, P., Huang, C.M., Li, C.J., Vagbo, C.B., Shi, Y., Wang, W.L., Song, S.H., Lu, Z., Bosmans, R.P., Dai, Q., Hao, Y.J., Yang, X., Zhao, W.M., Tong, W.M., Wang, X.J., Bogdan, F., Furu, K., Fu, Y., Jia, G., Zhao, X., Liu, J., Krokan, H.E., Klungland, A., Yang, Y.G., and He, C. (2013). ALKBH5 is a mammalian RNA demethylase that impacts RNA metabolism and mouse fertility. Molecular cell 49, 18–29. https://doi.org/10.1016/j.molcel.2012.10.015

Zhou, H., Su, J., Hu, X., Zhou, C., Li, H., Chen, Z., Xiao, Q., Wang, B., Wu, W., Sun, Y., Zhou, Y., Tang, C., Liu, F., Wang, L., Feng, C., Liu, M., Li, S., Zhang, Y., Xu, H., Yao, H., Shi, L., and Yang, H. (2020). Glia-to-Neuron Conversion by CRISPR-CasRx Alleviates Symptoms of Neurological Disease in Mice. Cell 181, 590–603 e516. https://doi.org/S0092-8674(20)30286-5

Zuchero, J.B., and Barres, B.A. (2015). Glia in mammalian development and disease. Development (Cambridge, England) 142, 3805–3809. https://doi.org/10.1242/dev.129304

